# Different methods for niche and fitness differences computation offer contrasting explanations of species coexistence

**DOI:** 10.1101/2021.09.28.462166

**Authors:** Jurg W. Spaak, Po-Ju Ke, Andrew D. Letten, Frederik De Laender

**Author notes:** Corresponding author, phone-number. Statement of authorship:* J.W.S. wrote the computer code. J.W.S. wrote the first draft, all authors discussed content and contributed to revisions. Data accessibility:* All computer code used in this manuscript will be made publicly available on figshare.

## Abstract

In modern coexistence theory, species coexistence can either arise via stabilizing mechanisms that increase niche differences or equalizing mechanisms that reduce fitness differences. Having a common currency for interpreting these mechanisms is essential for synthesizing knowledge across different studies and systems. Several methods for quantifying niche and fitness differences exist, but it remains unknown to what extent these methods agree on the reasons why species coexist. Here, we apply four common methods to quantify niche and fitness differences to one simulated and two empirical data sets. We ask if different methods result in different insights into what drives species coexistence. We find that different methods disagree on the effects of resource supply rates (simulated data), and of plant traits or phylogenetic distance (empirical data), on niche and fitness differences. More specifically, these methods often do not agree better than expected by chance. We argue for (1) a better understanding of what connects and sets apart different methods, and (2) the simultaneous application of multiple methods to enhance a more complete insight into why species coexist.

## Introduction

Explaining biodiversity is a central goal in ecology (Hutchinson, 1959). There are many different perspectives to address this objective. Neutral theory focuses on regional processes (Hubbell & Knapp, 2003), contemporary niche theory focuses on investigating species’ limiting factors (Tilman, 1982), and modern coexistence theory focuses on separating niche and fitness differences that help and hamper coexistence (Adler *et al*., 2007).

Ecologically equivalent species can’t coexist, as stated by the competitive exclusion principle (Levine & HilleRisLambers, 2009). To stably coexist, species must differ in some aspect. For example, if species differ in their resource uptake traits, this difference may contribute to negative frequency dependence and help rescue a rare species from competitive exclusion (Letten *et al*., 2017). Under modern coexistence theory, such differences are termed niche differences (Spaak *et al*., 2020; Chesson, 2000), which capture all species differences that help a species to recover from low density. Mechanisms that increase niche differences are called stabilizing mechanisms. Examples include resource partitioning, different root lengths, or species-specific predators. However, species differences may not only increase niche differences, but can also increase competitive differences. For example, differences in rooting length of plant species may not only cause a species to consume resources at different depth, but also affect how much resources a species can take up (Adler *et al*., 2007). Such differences are termed fitness differences, which capture all differences of species that are associated with competitive dominance. More precisely, fitness differences identify all differences in species performance in the hypothetical case that there are no niche differences. Mechanisms that decreases fitness differences are called equalizing mechanisms.

Modern coexistence theory assesses coexistence by computing invasion growth rates (Chesson, 2003; Barabás *et al*., 2018; Ellner *et al*., 2018; Schreiber, 2000). Niche and fitness differences, which are derived from the decomposition of invasion growth rates, are tools that help us understand the mechanisms underlying species coexistence (Spaak *et al*., 2021; Carmel *et al*., 2017; Carroll *et al*., 2011; Adler *et al*., 2007; Song *et al*., 2019). While the invasion growth rates often suffice to predict coexistence, niche and fitness differences can in principle offer additional understanding of why the species do or do not coexist (Grainger *et al*., 2019; Ke & Letten, 2018; Spaak *et al*., 2020, 2021). Without such additional understanding the effort of computing niche and fitness differences is unnecessary.

Unfortunately, computing niche and fitness differences is not a straightforward task. There are many different methods to compute niche and fitness differences (Spaak *et al*., 2020), with new methods still being developed (Johnson, 2021; Koffel *et al*., 2021). Some of these methods are tailored to a specific community model (Godoy & Levine, 2014; Saavedra *et al*., 2017; Chesson, 2018), while others are model agnostic (Spaak *et al*., 2020; Carroll *et al*., 2011; Zhao *et al*., 2016). Few of these methods have been applied extensively to experiments, while others have been used only few times and little is known about their practical usability (Adler *et al*., 2007; Bimler *et al*., 2018; Zhao *et al*., 2016; Spaak *et al*., 2020; Carmel *et al*., 2017). Despite the proliferation of different methods to assess niche and fitness differences, there is very little guidance on the insights offered by different methods on the mechanisms underpinning coexistence (Spaak *et al*., 2020, 2021; Godwin *et al*., 2020).

Optimally, all these different methods to compute niche and fitness differences would lead to the same insights. That is, independent of which method we apply to our data, we would come to the same conclusion regarding, for instance, which traits increase niche or fitness differences (Kraft *et al*., 2015; Pérez-Ramos *et al*., 2019), how phylogenetic distance affects niche and fitness differences (Narwani *et al*., 2013; Germain *et al*., 2016), or how the presence of trophic interactions increases fitness differences (Petry *et al*., 2018; Terry & Lewis, 2020).

However, here we show that this is not always the case. We do so by analysing three data sets (Letten *et al*., 2017; Pérez-Ramos *et al*., 2019; Germain *et al*., 2016), using four different methods to compute niche and fitness differences. This exercise shows that different methods can lead to different conclusions regarding the putative mechanisms driving coexistence. The first data set consisted of simulation output from a virtual experiment in which we investigated how changes in resource availability affect niche and fitness differences, similar to the work performed by Letten *et al*. (2017). We find that the different methods often do not agree on how resource availability affects niche and fitness differences. The second data set contains empirically measured interaction strengths among, and traits of, annual plant species. We asked which traits best predict niche and fitness differences, similar to the work of Pérez-Ramos *et al*. (2019). We find that different methods lead to different conclusions on which traits best predict niche and fitness differences. The third data set contains empirically measured interaction strengths among, and phylogeny information of, annual plants. We assess under which conditions phylogenetic distances hold information about niche or fitness differences, similar to the work of Germain *et al*. (2016). Again, we find that the method used to assess niche and fitness difference influences the answer to this question.

## Methods

### Methods to assess niche and fitness differences

There are as many as 13 different methods to compute niche and fitness differences (Spaak *et al*., 2020; Koffel *et al*., 2021; Johnson, 2021). In the current work, we focus on those methods that are most widely applied and/or are applicable to most data. The original definition of the four selected methods can be found in Chesson & Kuang (2008); Carroll *et al*. (2011); Zhao *et al*. (2016) and Spaak *et al*. (2020). Specifically, we chose these four methods as they are the only ones that can be applied to all three investigated datasets (see below).

For better comparison between the four methods we have slightly adjusted them. We changed the definitions such that a community with no niche differentiation will indeed have zero niche differences and increasing the stabilizing mechanisms will increase niche differences. We changed the definition of fitness differences such that the two neutral species have fitness differences of 1. Here, we assigned a fitness difference to each species, i.e. ℱ_*i*_, and the species with fitness difference below 1 is the competitive superior species, which would competitively exclude the other species in the absence of niche differences. In all cases this was achieved by a linear transformation of the original definitions, which does not affect the underlying conceptual ideas of the methods.

The first and original method to compute niche and fitness differences is based on a Lotka-Volterra model, which is derived from an underlying MacArthur consumer-resource model with the potential inclusion of a higher trophic level (Chesson, 1990; Chesson & Kuang, 2008; Chesson, 2013).

We assume a two-species Lotka-Volterra model 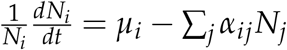, where *N*_*i*_ is the density of species *i, µ*_*i*_ is the intrinsic growth rate of species *i* and *α*_*ij*_ is the per-capita effect of species *j* on species *i*. For this model, the niche and fitness differences are defined as

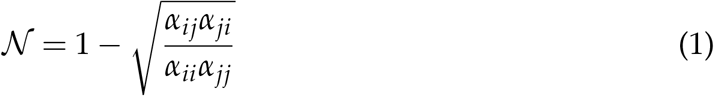

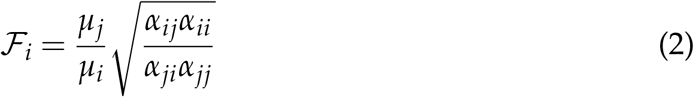

Usually, niche differences are described as 1 − *ρ* in the literature, where 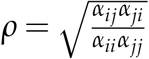 is the niche overlap. We did not alter the definition of this method (Godoy & Levine, 2014; Letten *et al*., 2017; Chesson, 2013). This method is originally only defined for a two species Lotka-Volterra model, however, it can be applied to other community models by reducing a given community model to Lotka-Volterra form. Specifically, Letten *et al*. (2017) have adapted this method to the Tilman consumer-resource model and Godoy & Levine (2014) have adapted this method to the Beverton-Holt annual plant model. We effectively use their computations of niche and fitness differences for the examples below. We will refer to this method as the *Original* method to assess niche and fitness differences.

The first method that we have introduced is model specific, all other methods can be applied to a larger range of community models and can all be interpreted as a decomposition of the invasion growth rates. We therefore first introduce the notion of invasion growth rates. We assume a community model of the form 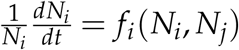 We assume that both species have a stable monoculture equilibrium density, denoted 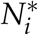 and 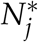, respectively. The invasion growth rate of species *i* is the growth rate when it is at low density (mathematically zero) and its competitor at their monoculture density, i.e. the invasion growth rate is denoted as 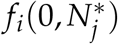. Another important growth rate is the intrinsic growth rate when both species are at low density, i.e. *f*_*i*_(0, 0).

The second method, as defined by Carroll *et al*. (2011), uses the sensitivity of each species to competition, i.e. 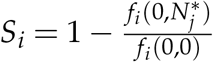. With this they define

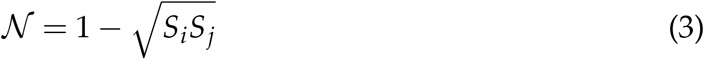

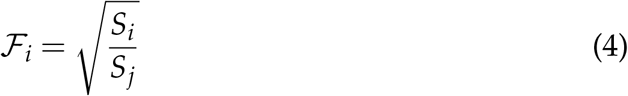

The original work uses the notation *ND* and *RFD* for niche and fitness differences, respectively. The only difference is that *RFD* in the original studies corresponds to the maximum of ℱ_*i*_ and ℱ_*j*_. Importantly, this definition yields identical results for the Lotka-Volterra model as the original method. However, it is not equivalent to the derivations of Godoy & Levine (2014) or Letten *et al*. (2017) for their respective models. We will refer to this method as the *geometrical* method, because it computes niche and fitness differences as the geometrical mean and variance of the sensitivities, *S*_*i*_.

The third method, by Zhao *et al*. (2016), define niche and fitness differences as

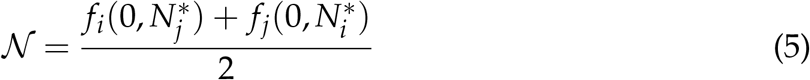

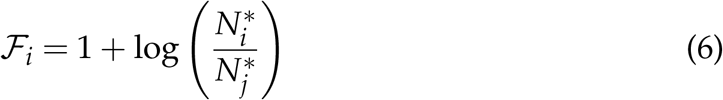

We have adjusted their definition in the following way 𝒩 = (−*ρ*)/2. where *ρ* is their definition of niche overlap and 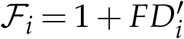, where 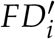 is the original definition by Zhao *et al*. (2016); this was done to ensure that a neutral community would indeed have zero niche differences. The method by Zhao *et al*. (2016) is not equivalent to the original method for the Lotka-Volterra model. We will refer to this method as the *arithmetical* method as they compute niche differences as the arithmetic mean of the invasion growth rates.

The last method we investigate is the method by Spaak *et al*. (2020), who defined 𝒩 and ℱ as

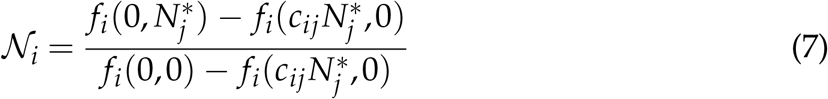

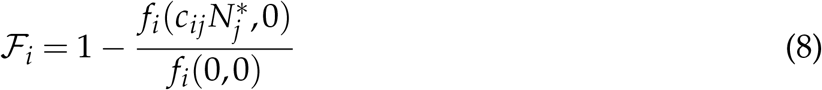

Where *c*_*ij*_ is a conversion factor that converts densities of species *j* to densities of species *i*, while keeping the converted density constant. *c*_*ij*_ is the solution of the equation |1 − 𝒩_*i*_| = |1 − 𝒩_*j*_|. The current work does not include any facilitation, i.e. 𝒩_*i*_ > 1, which implies 𝒩_*i*_ = 𝒩_*j*_, we therefore drop the subscript *i*. We have adjusted their definition in the following way, 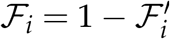, where 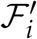 is the original definition by Spaak *et al*. (2020).

Equation 7 compares the actual invasion growth rate of species 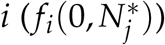 to the hypothetical invasion growth rate when the two species would not interact (*f*_*i*_(0, 0)) and to another hypothetical invasion growth rate when the two species had no niche differences 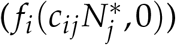. Importantly, this definition yields identical results for the Lotka-Volterra model as the original method, similar to the method by Carroll *et al*. (2011). We will refer to this method as the *species-specific* method, as niche and fitness differences are species-specific properties and not community specific properties in this method. For a more in detail explanation of all these methods we refer to the original papers (Carroll *et al*., 2011; Zhao *et al*., 2016; Spaak *et al*., 2020).

### Examples

We have chosen three studies from the literature, and performed similar analysis of their data, but with four instead of only one definition for niche and fitness differences. We have chosen these three studies because they belong to the core literature of modern coexistence theory. As such, we assume and hope that many readers will be familiar with their work. Importantly, we do not re-analyze the data of those three studies identically as described in the original study, but simplified (e.g., random effect structure, error distributions). As such, our results cannot be directly compared with the original material even if using the same method, but rather, can be used to compare how interpretations may differ using different estimates of niche and fitness differences. We had no a-priori expectations on the robustness of the results of these three studies, and we do not want to imply that our reevaluation of their results should cast doubt on them. In fact, the three studies are all more complex and insightful than presented here, we invite any reader not familiar with their work to read the original papers.

### Theoretical example

Letten *et al*. (2017) investigated how changes of parameters in a two-species consumer-resource model affect niche and fitness differences. They used a Tilman consumer-resource model with substitutable resources with a Holling type 2 response. The community dynamics for the substitutable resources are given by

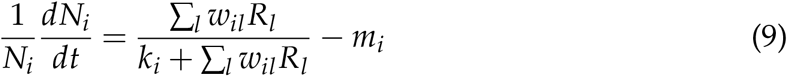

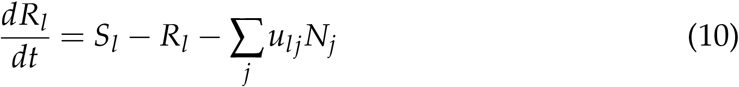

where *w*_*il*_ is the conversion of resource *l* to biomass of species *i, u*_*li*_ is the utilization of resource *l* by species *i, m*_*i*_ is the mortality rate, *S*_*l*_ is the resource supply and *k*_*i*_ is half-saturation constant.

Letten *et al*. (2017) investigated how niche and fitness differences change in response to a change in all of the parameters (*u*_*li*_, *w*_*il*_, *k*_*i*_, *S*_*l*_). We will, however, only investigate how niche and fitness differences change as a function of *S*_*l*_. Specifically, we assess how a change in resource supply rates *S*_*l*_ increases or decreases niche and fitness differences.

Additionally, we investigate whether the different methods agree on which species is the superior competitor and whether the different methods agree on the frequency dependence of the system. The superior competitor is defined as the species with ℱ_*i*_ < 1. The frequency dependence of the system is defined by the sign of the niche differences 𝒩: negative niche differences correspond to *positive* frequency dependence and positive niche differences correspond to *negative* frequency dependence. 𝒩 = 0 indicates the absence of frequency dependence.

### Empirical examples

Many different papers have tried to identify predictors of niche and fitness differences, such as phylogenetic distance (Narwani *et al*., 2013; Germain *et al*., 2016), the presence of predators (Chesson & Kuang, 2008; Petry *et al*., 2018; Terry & Lewis, 2020) or species traits (Pérez-Ramos *et al*., 2019; Kraft *et al*., 2015; Gallego *et al*., 2019). We chose to perform an analysis similar to that in Pérez-Ramos *et al*. (2019) (i.e., which trait differences are most important for predicting niche and fitness differences) and Germain *et al*. (2016) (i.e., under what circumstances does phylogenetic distance predict niche and fitness differences). We chose Pérez-Ramos *et al*. (2019) and Germain *et al*. (2016) as they offer a large and open data set with many species and many predictors of niche and fitness differences.

Pérez-Ramos *et al*. (2019) investigated whether differences in species traits translate to niche or fitness differences. Pérez-Ramos *et al*. (2019) parameterized pairwise competition models for 10 annual plant species (a total of 45 two-species annual plant communities). Additionally, Pérez-Ramos *et al*. (2019) measured functional traits for each species and correlated species’ differences in traits to their niche and fitness differences.

Germain *et al*. (2016) investigated under which circumstance phylogenetic distance can be used as a predictor of niche or fitness differences. Germain *et al*. (2016) investigated 30 annual plants and fitted an annual plant model to 20 two-species annual plant communities under two different environmental conditions (wet versus dry), leading to a total of 40 annual plant community models. Half of the species pairs were found in similar habitats (sympatric) and half were found in different habitats (allopatric). Additionally, they measured the phylogenetic distance of the two competing species to assess under which conditions phylogenetic distance affects niche and fitness differences.

With each study’s fully-parameterized annual plant model, we compute niche and fitness differences as defined by the original method using equations 1 and 2 and simulated the necessary growth rates for the other methods. We then perform a linear regression of the predictor (trait differences or phylogenetic distance between the two species) and the response variables, niche or fitness differences. For each linear regression we report the slope and the p-value.

## Results

### Theoretical example

First, we investigate two different communities of two species competing for two substitutable resources (Fig. 1, A,C). For each community, we changed the resource supply rates (from black triangle to black dot).

**Figure 1:**
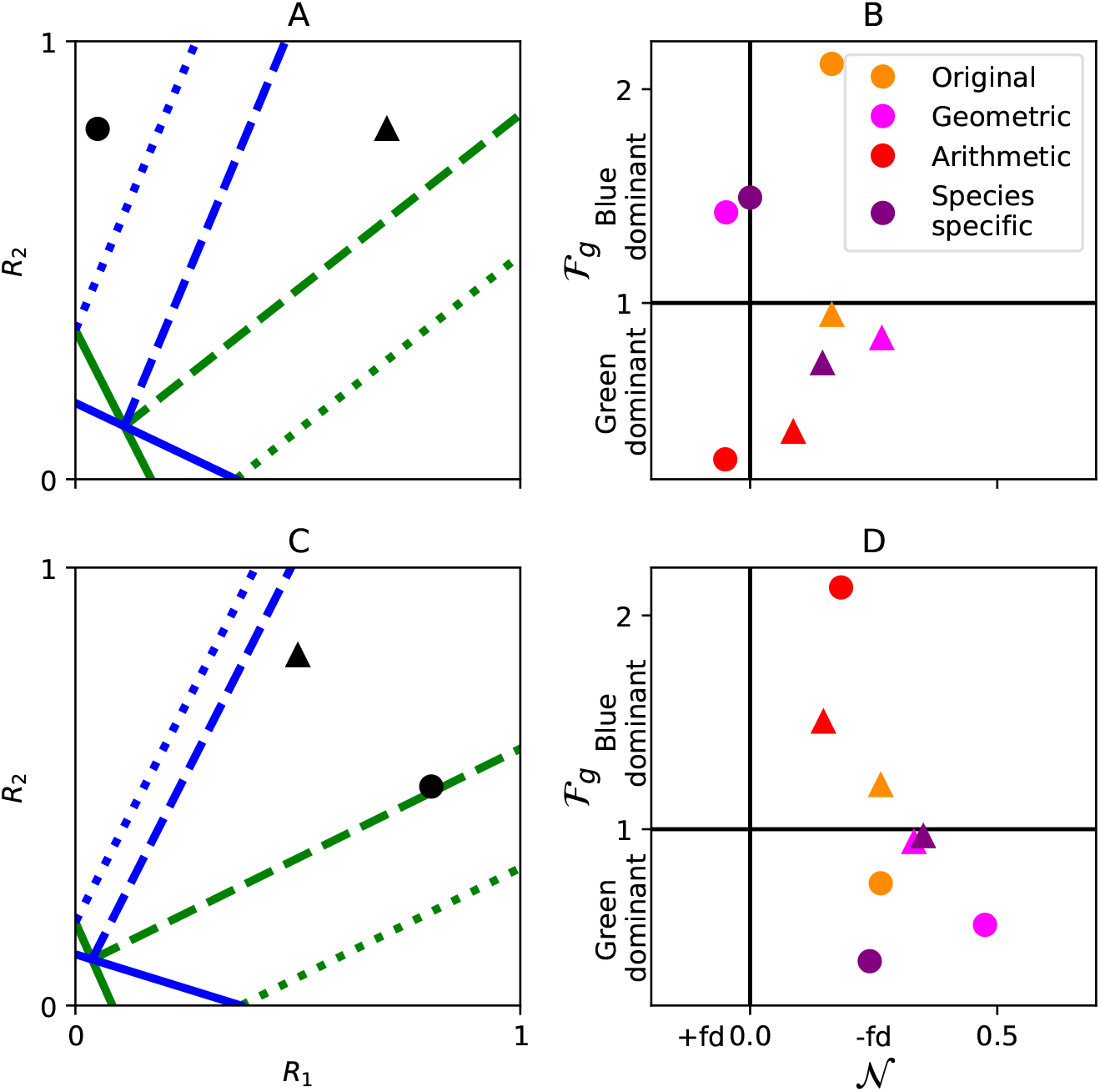
A,C: Two examples of two species competing for substitutional resources. The solid lines indicate the zero-net growth isoclines (ZNGI), at which the species has zero growth. The dashed lines indicate the consumption vectors *u*_*li*_ of the species. The black triangle and dot show the resource supply rates before and after an environmental change. B,D: Corresponding niche and fitness differences of the green species for the four investigated methods (colors). In certain cases, the different methods do not agree qualitatively on whether: 1. the community is driven by positive (𝒩 < 0, +fd), negative (𝒩 > 0 +fd) or no frequency dependence (Panel B, dot). 2. the blue species (ℱ_*g*_ > 1) or the green species (ℱ_*g*_ < 1) is the competitive superior species (Panel D, triangle). 3. the change in resource supply rates is more beneficial for the blue (higher ℱ_*g*_ for the dot than for the triangle) or green species (lower ℱ_*g*_ for the dot than for the triangle, Panel B); 4. the change in resource supply rates increases, decreases or has no effect on niche differences (Panel D, change in 𝒩) This figure is conceptually similar to Figure 2 from Letten *et al*. (2017). For a more quantitative assessment of these cases see Figure 2.

As observed before (Spaak *et al*., 2020; Song *et al*., 2019), the different methods have different quantitative values for 𝒩 and ℱ. However, in addition to differences in their quantitative values, the different methods also yield qualitatively different interpretations of the change of resource supply rates.

For example, in the first community (Panels A,B), we might ask whether the decrease in resource 1 is most beneficial for the blue or the green species. If the increase of resource 1 benefits the blue species, consistent with resource ratio theory (Tilman, 1982), then fitness differences of the green species, ℱ_*g*_, will increase. Conversely, ℱ_*g*_ will decrease if it benefits the green species. Yet, the different methods give different results. The original, geometric and species-specific methods all consider the change in resource 1 as more beneficial for the blue species. In contrast, the arithmetic method interprets this change as most beneficial for the green species.

Alternatively, we might ask whether the community is driven by positive or negative frequency dependence, which would lead to negative or positive niche differences, respectively. Again, we find that the different methods do not always lead to the same conclusion. At the new resource supply point (black dot), the original method finds that the community is driven by negative frequency dependence, as niche differences are positive. On the other hand, the geometric and arithmetic method interpret that the community is driven by positive frequency dependence. Finally, the species-specific method interprets that frequency dependence is exactly zero.

In the next community (Panels C,D), we ask about the stabilizing effect of resource change. The original method finds that the change in resource supply does not affect niche differences (Letten *et al*., 2017). The arithmetic and geometric method find that the change is stabilizing (increase in 𝒩), while the species-specific method interprets the change as destabilizing (decrease in 𝒩). Alternatively, we might ask which species is the competitive superior. Again, we obtain different answers when using different methods. The original and arithmetic method conclude that the blue species is the competitive superior (ℱ_*g*_ > 1), while the geometric and the species-specific methods conclude that the green species is the competitive superior (ℱ_*g*_ < 1, albeit with close to no fitness difference).

Admittedly, we have selected communities to illustrate our message that different methods can give different explanations as to why species coexist. We will now show that these are not (rare) special cases and that disagreement among methods is a persistent feature. To this end, we randomly generated 1000 two-species communities competing for substitutable resources (see Appendix S1) and computed their niche and fitness differences (Figure 2A). We then checked whether the methods agree qualitatively on frequency dependence (𝒩 > 0 or 𝒩 < 0) and the identity of the competitive dominant species (ℱ_*i*_ > 1 or ℱ_*i*_ < 1) of the community at the initial resource supply rate. This quantitative analysis confirms our qualitative examples; often, the methods did not agree well (red dots, Figure 2A) and they did not agree much better than expected by chance (Appendix S1).

**Figure 2:**
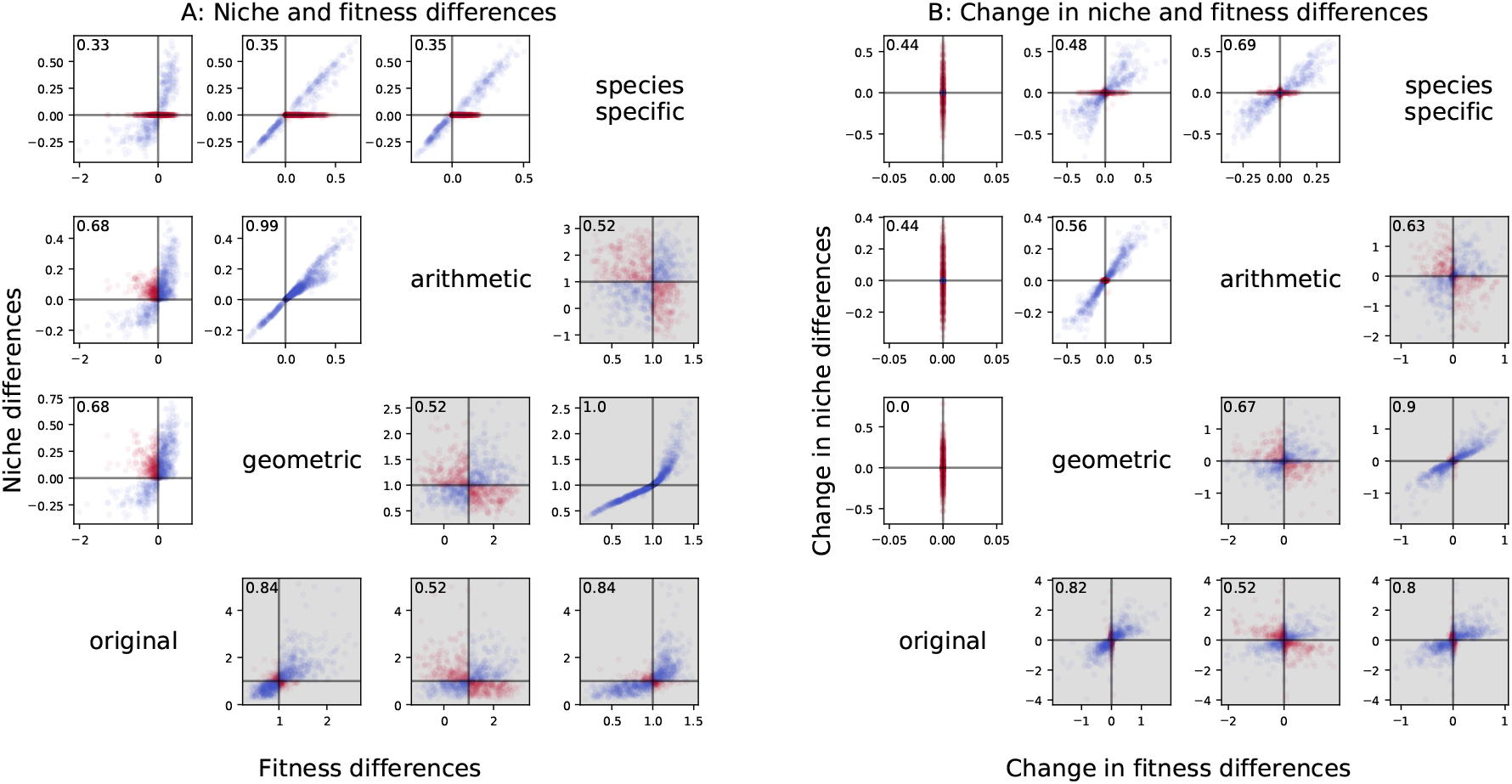
A: For 1000 randomly generated communities we computed niche (above diagonal, white panels) and fitness differences (below diagonal, grey panels) according to the four investigated methods (diagonal). Each dot corresponds to one randomly generated community. We say that two methods agree qualitatively if they agree on the frequency dependence (𝒩 > 0 or 𝒩< 0) and superior competitor (ℱ_*i*_ > 1 or ℱ_*i*_ < 1). Communities where the methods agree qualitatively are shown in blue, communities where the methods disagree are shown in red. The top-left number indicates how often the two methods agree qualitatively (fraction of blue dots). B: Similar to A, but we compare the change of niche and fitness differences after a change in resource supply rates. We say that two methods agree qualitatively when niche or fitness differences increase or decrease for both methods. A,B: In general the different methods are well correlated. However, their qualitative agreement (top left number = fraction of blue points) is often not much higher than expected by chance (see Appendix S1).

Next, we changed the resource supply rates and computed niche and fitness differences for the changed resource supply rate. We assessed whether the methods agree on the stabilizing effect (increase or decrease in 𝒩) and on the equalizing effect (increase or decrease in ℱ_*i*_) of the change in resource supply rate (Figure 2B). Again, the different methods did not agree well on how changes in resource supply rates affect niche and fitness differences.

The only case with 100% agreement was the one where the geometric and the species-specific method identify the superior competitor, i.e. ℱ_*i*_ > 0 or ℱ_*i*_ < 0. We show in the appendix why this is always the case (Appendix S3). For all other cases, the specific percentages depend on the specifics of the randomly generated communities, yet the qualitative picture, that the methods do not always agree is very consistent (Appendix S1). Changes in resource supply rates do not affect niche differences when the original method was applied to the model in equation 9 and 10 (Letten *et al*., 2017), whereas it can affect niche differences for the other methods.

### Empirical examples

An important empirical application of modern coexistence theory is to link traits to niche and fitness differences. Doing so is important to provide insight into which processes govern coexistence in nature (Kraft *et al*., 2015; Gallego *et al*., 2019; Germain *et al*., 2016; Narwani *et al*., 2013; Godoy & Levine, 2014). To do so, one typically measures the relationship between traits and niche and fitness differences. For example, if we hypothesize that species with different root length will consume resources at different locations (Levine & HilleRisLambers, 2009), we can test the correlation between differences in root length and niche differences. Alternatively, we might hypothesize that species with longer roots will have a competitive advantage, as they can take up more resources, which can be tested by quantifying the correlation between species differences in root length and fitness differences.

We computed niche and fitness differences for competing annual plant species with the four methods mentioned above. We then tested which trait differences were correlated to niche and fitness differences (Fig. 3). The different methods yielded different quantitative and qualitative results about the importance of different traits.

**Figure 3:**
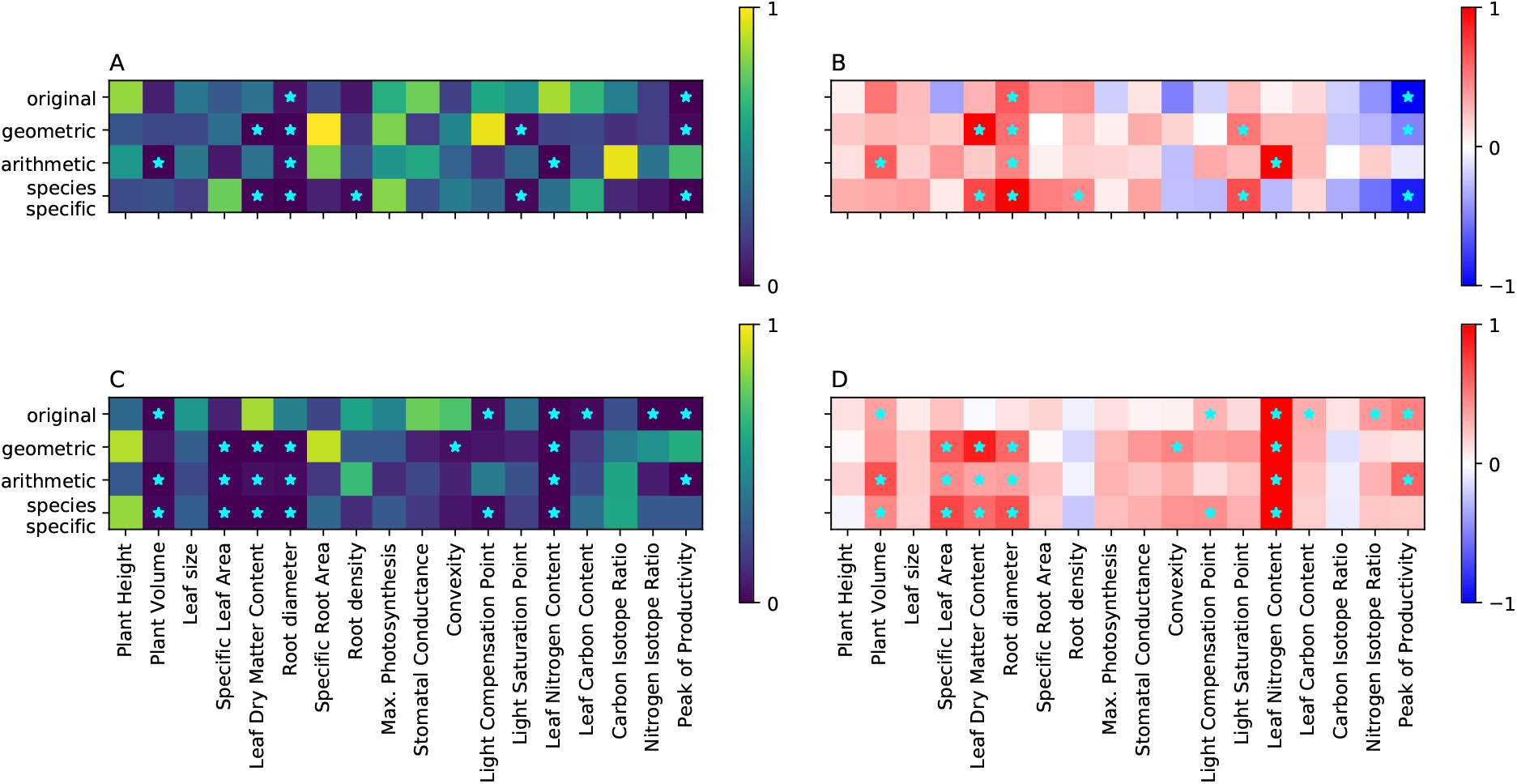
The p-value (A,C) and the scaled slope (B,D) of the linear regression between the pairwise trait differences (x-axis) and the niche (A,B) and fitness differences (C,D) for each of the four investigated methods (y-axis) for the Pérez-Ramos *et al*. (2019) data set. We find many quantitative and qualitative differences among these linear regressions. For example, the original method predicts that leaf carbon content is a good predictor of fitness differences (cyan-star indicates p-value < 0.05) while none of the other methods do. Whether the experiments would have been analyzed by the original method or not will therefore affect our understanding of the system. We re-scaled the slopes of the linear regression for illustration purposes, for each method. The slopes were scaled such that the absolute value of the maximal slope is 1. Importantly, this does not qualitatively alter the slopes, as their sign and relative importance are not altered.

Quantitatively, different methods predicted different p-values (Fig. 3 A,C) and different slopes (Fig. 3 B,D) of the relationships between traits and niche and fitness differences. However, these differences are also qualitative and offer different answers to the question which traits drive coexistence. For example, only the geometric and species-specific method find differences in leaf dry matter content as significant predictors for niche differences. Similarly, only the original and arithmetic method find differences in peak productivity as significant predictors for fitness differences. Additionally, sometimes the slopes between the different methods are different, as can be seen by the example of the specific leaf area for niche differences. However, in the investigated dataset, we never observed two methods predicting qualitatively different slopes that were both significant.

In our third example we investigate in which scenarios phylogenetic distance is a good predictor of niche or fitness differences. Germain *et al*. (2016) conducted experiments with four different scenarios in a two-by-two factorial design: whether the species evolved under sympatry or allopatry, and whether the environment was wet or dry.

Again, we find that the different methods yield different results. For example, phylogenetic distance of species is a good predictor of niche and fitness differences for allopatrically-evolved species, when these two components are as measured by the original and arithmetic method, but not by the others. Conversely, phylogenetic distance of species that evolved sympatrically is a good predictor of fitness differences only for the geometrical and species-specific method.

For the two empirical datasets investigated here, we computed how often they agree qualitatively. That is, how often do two different methods give the same prediction of whether a given trait or phylogenetic distance significantly predicts niche or fitness differences (Fig.5). On the one hand, the different methods agree fairly well (agreement ranging from 55% − 82% for niche differences and 65% − 96% for fitness differences). On the other hand, most of the methods do not agree much better than expected by chance (60% − 65% and 50% − 62% for fitness differences). In general, the methods seem to agree more for fitness differences than for niche differences. The geometric and the species-specific methods agree best in general.

## Discussion

We have compared four different methods to compute niche and fitness differences on one theoretical and two empirical datasets. We found that the different methods can lead to different qualitative insights about the underlying ecology of the system. For example, the methods did not agree on whether a change in resource supply rates is stabilizing or destabilizing. Similarly, they did not agree on which traits are most important for niche and fitness differences. We have shown that these differences are not special cases, but rather the norm. We found that the different methods agree only slightly better than expected by chance (Fig. 2 and 5).

It is well known that these two key concepts of modern coexistence theory (niche and fitness difference) are not defined universally. The presented results show that this variation can lead to different interpretations, confirming results by Song *et al*. (2019). These authors showed that stabilizing and equalizing mechanisms have two different meanings in modern coexistence theory, depending on whether one investigates a two-species community or a multi-species community. We have generalized their work and showed that even for simple two species communities different methods can produce different assessments of stabilizing and equalizing mechanisms. There are other methods to assess niche and fitness differences which we have not examined. We expect that including other methods would increase this variation as different methods have different requirements and capture different aspects of coexistence.

### Perspectives on quantifying niche and fitness differences

Intuitively, the different methods lead to different interpretations of coexistence because they capture different aspects of ecology. This is mirrored by which information is included into the calculation of niche differences (Appendix S3). The original method includes inter- and intra-specific species interactions; the arithmetic method includes invasion growth rates; the geometric method includes invasion and intrinsic growth rates; and the species-specific method includes invasion, intrinsic and no-niche growth rates. These conceptual differences show that different information is included in the computation of niche and fitness differences, which lead to qualitatively and quantitatively different interpretations of coexistence.

For example, a change in resource supply rates that increases invasion growth rates, e.g. increasing resource supply rates at fixed ratios, will increase niche differences according to the arithmetic method. However, this change in resource supply rates will likely also affect the other growth rates, e.g. increase intrinsic growth rate and decrease no-niche growth rate (Appendix S2). Whether niche differences according to the geometric and species-specific method also increase depends on the magnitude and direction of the change in these growth rates. How different methods interpret changes in resource supply rates therefore mirror how the intrinsic, invasion and no-niche growth rates are affected by these resource supply rate changes, as well as whether this information is included. A similar understanding of the empirical data-sets can explain how different methods interpret different traits or phylogenetic distances as predictors of niche and fitness differences (Appendix S2).

We found that different methods yield different niche and fitness differences, and different values for stabilizing and equalizing mechanisms. We argue that identifying which method is the superior one is both impossible and unwarranted. Instead, we feel that the different methods considered merely reflect different interpretations of these notoriously slippery concepts. The objective is therefore not to seek hierarchy, but understanding across methods, much like what is done in other research fields. Indeed, variation of interpretation, and the resulting methodological variation, is found back in other research fields as well. For instance, concepts like stability and biodiversity can be translated in various ways into mathematical formulae, depending on what one considers a “stable” or “diverse” ecosystem, or which aspects one wishes to highlight (Montoya *et al*., 2018; Donohue *et al*., 2016; Mcgill *et al*., 2015). In stability research, this variation has been embraced and coined the dimensionality of stability (Radchuk *et al*., 2019; Donohue *et al*., 2016). Understanding why these methods differ and when they correspond, i.e. when dimensionality is reducible, is a major objective in stability research (Radchuk *et al*., 2019; Arnoldi *et al*., 2018; Clark *et al*., 2021; Carpentier *et al*., 2021). We argue that similar efforts are needed in modern coexistence theory, if we want to gain synthetic understanding across published results adopting a variety of methods.

Additionally, the best-suited method to quantify certain aspects of niche and fitness differences may depend on the particular empirical system Godwin *et al*. (2020). Of the methods examined here, some require parameterizing a demographic model (Chesson & Kuang, 2008; Chesson, 2013) whereas others require measuring species’ invasion growth rate (Carroll *et al*., 2011; Zhao *et al*., 2016; Carmel *et al*., 2017). While the latter methods are potentially model-independent, they are only better suitable when the monoculture equilibrium can be ensured and the invasion growth rate can be quantified within a feasible time frame; these criteria likely explain the prevalence of such methods when studying the coexistence of microorganisms (e.g. Narwani *et al*., 2013; Grainger *et al*., 2019; Li, 2017). On the other hand, quantifying niche and fitness differences for long-lived organisms, e.g. plants, were mainly achieved by parameterizing demographic models, e.g. Godoy & Levine (2014); Chu & Adler (2015). Although labour-intensive, we would argue that parameterizing a demographic model might be the most flexible approach when the goal is to assess niche and fitness differences with multiple methods. Indeed, we were able to calculate multiple metrics from the Pérez-Ramos *et al*. (2019) and Germain *et al*. (2016) data set precisely because they provided a fully-parameterized annual plant model, from which the original method can be calculated and invasion growth rates required by other methods can be simulated.

We therefore propose that future work in modern coexistence theory should consider applying multiple methods to assess niche and fitness differences. Often, one can with little to no additional effort compute multiple methods of niche and fitness differences with the same data (or with an underlying model). Computing multiple niche and fitness differences on the same data will help the individual researcher as well as the field of modern coexistence theory.

We see the two following benefits for the field of modern coexistence theory. First, we start to understand under which conditions the dimensionality of niche and fitness differences is high. This advances our understanding of when the different methods agree well and might increase our understanding of when intrinsic growth rates or no-niche growth rates are important for coexistence. Second, new methods to compute niche and fitness differences are continuously being developed (Spaak *et al*., 2020, 2021; Koffel *et al*., 2021; Johnson, 2021), however, these are rarely applied empirically. Applying multiple methods to one data set will allow us to understand the differences and potential benefits of newer methods earlier.

Additionally, each individual researcher benefits from applying multiple methods to their data. First, the different methods highlight different aspects of coexistence. The correctness of a certain hypothesis might depend on which part of coexistence is highlighted. By applying multiple methods, and therefore highlighting different aspects, one is more likely to highlight the correct aspect as well. Of course, this should not be misused for p-hacking or similar practices. Second, by highlighting different aspects and understanding under which scenario a hypothesis is correct one will also obtain a deeper understanding of coexistence and of one’s own data. For example, by which growth rate is the correctness of a hypothesis driven? Third, niche and fitness differences are difficult to grasp and each of the currently available methods describes them from a different perspective. By applying multiple methods we may get a better grasp of them. For example, phylogenetic distance in sympatric evolution is a good predictor of niche differences according to three different methods (Fig. 4 A). We can therefore be more certain that phylogenetic distance in sympatric evolution indeed affects niche differentiation.

**Figure 4:**
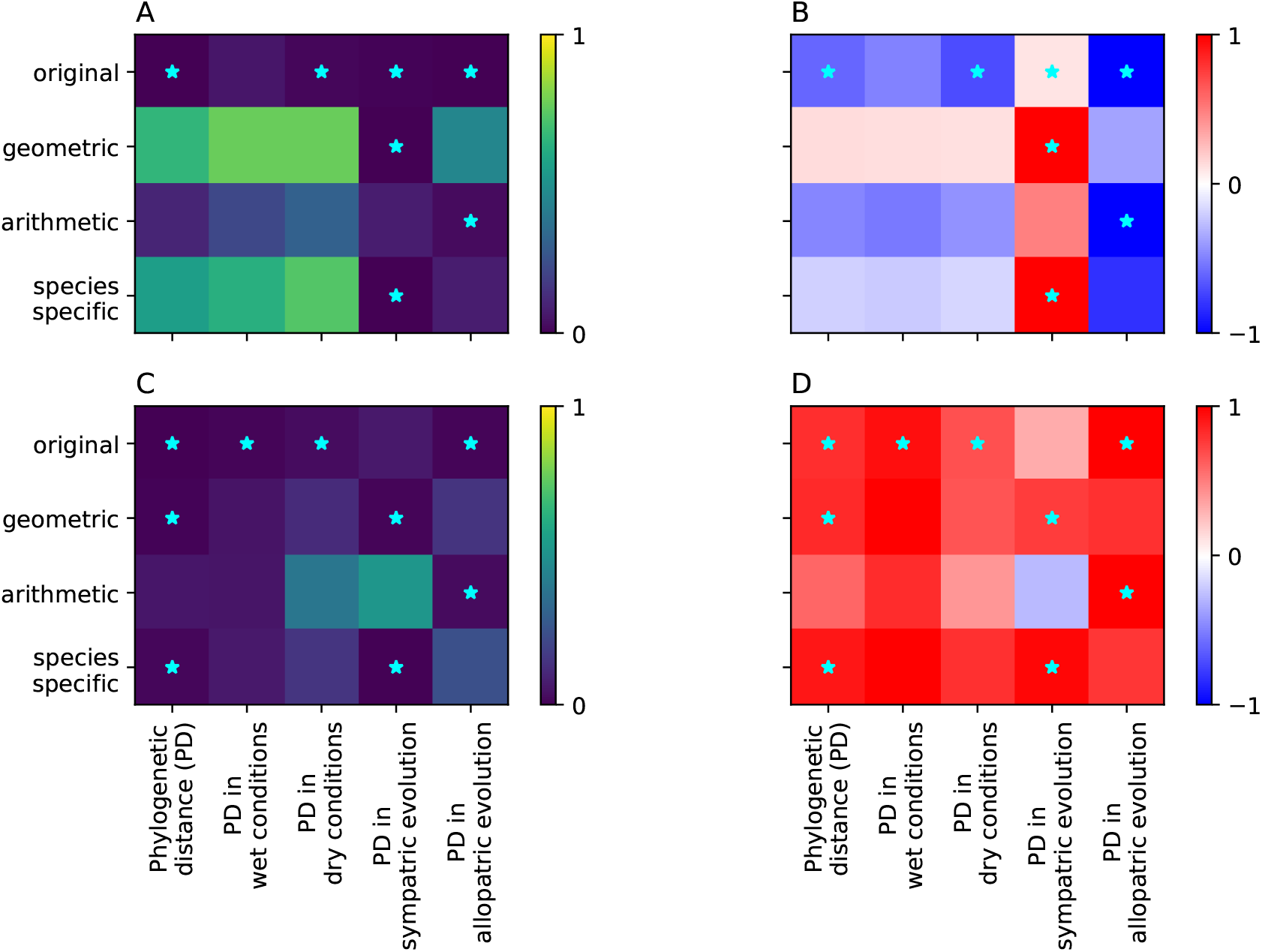
Similar to figure 3, we report the p-value (A,C) and the relative slope (B,D) of the linear regression between the pairwise phylogenetic distance (x-axis) and the niche (A,B) and fitness differences (C,D) for each of the four investigated methods (y-axis) for the Germain *et al*. (2016) data set. Again, we find that different methods lead to qualitatively different conclusions about in which scenario phylogenetic distance predicts niche or fitness differences. The first column (Phylogenetic distance; PD) reports the linear regression based on all communities, while the second and third column does so for communities in wet and dry conditions, respectively. The fourth and fifth column reports the results for sympatric and allopatric communities, respectively.

**Figure 5:**
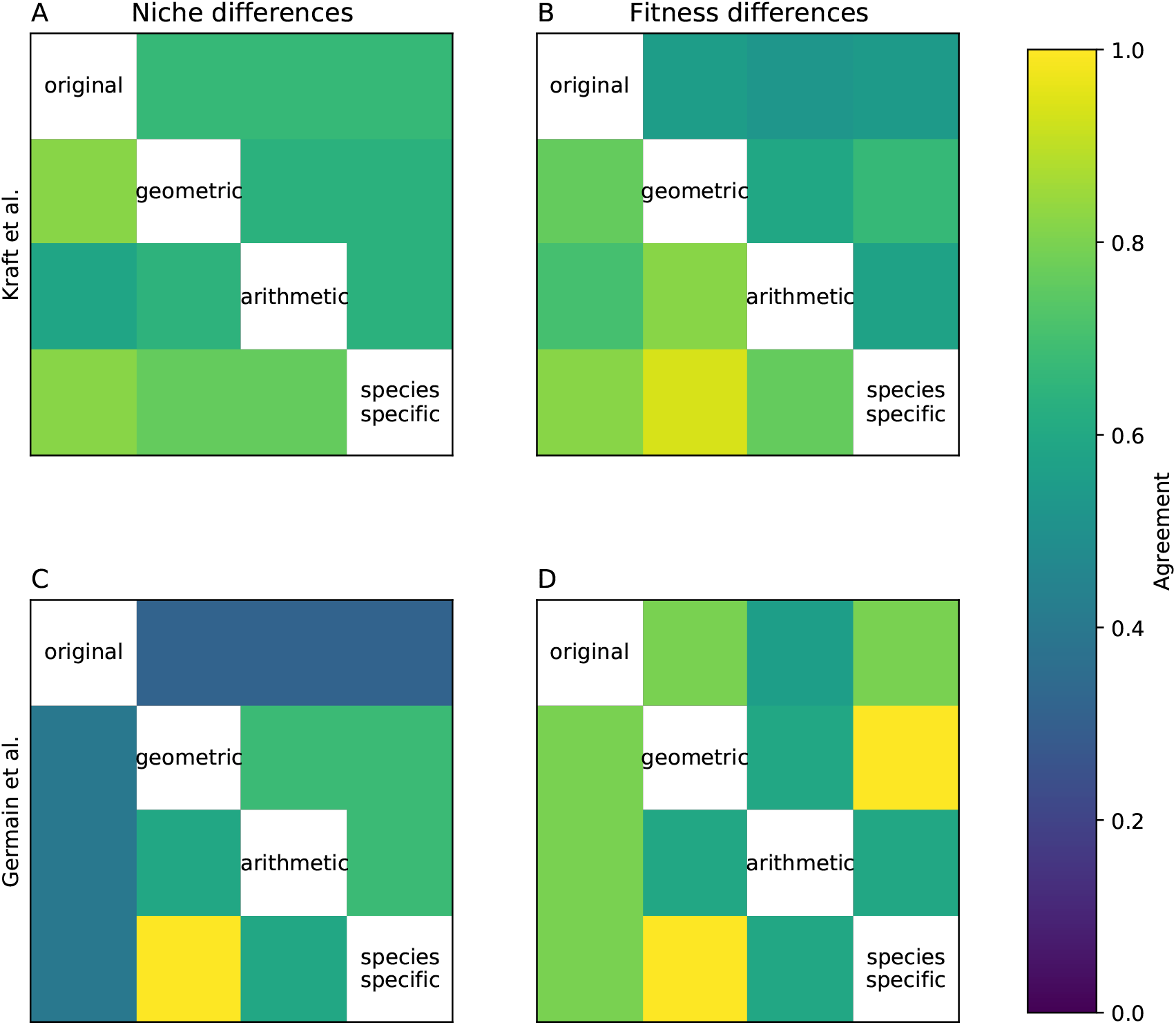
We report how often the two methods agree on whether a certain trait (A,B) or phylogenetic distance (C,D) is significant for niche (A,C) or fitness (B,D) differences. We compare how often they agree (below diagonal) to how often we would expect them to agree based on chance (above diagonal). While the methods, generally, agree fairly well, they do not agree much better than expected by chance.

## Acknowledgements

J.W.S received support from Schweizerischer Nationalfonds Early Post-doc mobility (P2SKP3 194960) F.D.L. received support from grant of the Fund for Scientific Research, FNRS (PDR T.0048.16). P.-J. K. received support from the Yushan scholar program of Taiwan MOE (NTU-110VV010). We thank Rachel Germain and Oscar Godoy for the shared data on the annual plant communities. We thank Rachel Germain and Oscar Godoy for comments on earlier versions of this manuscript.

## S1 Quantitative comparisons of methods for resource limitation

To create Figure 2 from the main text we created 1000 two-communities competing for two substitutional resources, given by the following per-capita growth rates

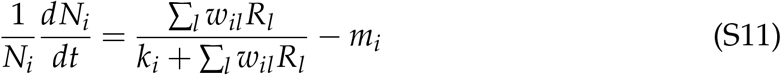

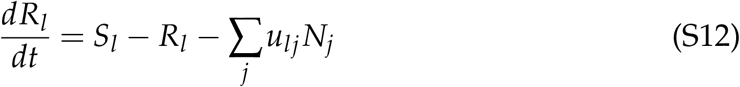

The species parameters were chosen randomly as given by table S1. For each two species community we computed niche and fitness differences in two different environmental settings. The two settings differed only in the resource supply rates *S*_*l*_, the distribution of the resource supply rates are also given in table S1. The parameter values were chosen randomly, there’s nothing special about the given parameter values and they do not necessarily represent realistic parameter values. We performed similar analysis with other parameter values and got qualitative similar results.

There are three qualitatively different effects the change of resource supply rates can have on niche or fitness differences: increase, decrease or have no effect. Figure 2 B show how often two different methods agree on the sign of the change in niche or fitness differences. To compute how often we would expect them to agree by chance we computed the probabilities *P*(*x*|*method*) where *x* is the sign of the change of niche or fitness differences (*x* ∈ {−1, 0, +1}) and method is any of the four investigated methods (Fig. S2). The base probability that the two methods agree is given by ∑_*x*∈{−1,0,1}_ *P*(*x*|*method*_1_)*P*(*x*|*method*_2_). For change in niche differences this probability was usually around 1/3, which is what we would expect naively, as there are 3 different possible outcomes. For change in fitness differences this probability was quite exactly 1/2. We never observed that a change in resource supply rates does not affect fitness differences, leaving effectively only two possible outcomes.

**Table S1:**
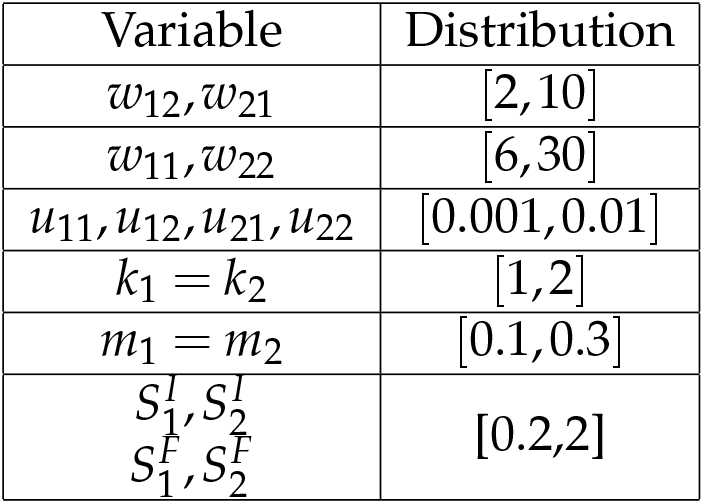
All distributions were assumed to be uniform distributions. We assumed that species 1 is a better competitor for resource 1 and species 2 is a better competitor for resource 2, we therefore chose different distributions for *w*_11_ and *w*_22_ than for *w*_12_ and *w*_21_. Additionally, we assumed that both species have identical mortality rate and identical half-saturation constants. This ensures that the zero net growth isoclines of the two species cross. All parameter were chosen arbitrarily. We investigated other parameter settings as well as competition for essential resources and got qualitatively similar results.

| Variable | Distribution |
| --- | --- |
| $w_{12}, w_{21}$ | $[2, 10]$ |
| $w_{11}, w_{22}$ | $[6, 30]$ |
| $u_{11}, u_{12}, u_{21}, u_{22}$ | $[0.001, 0.01]$ |
| $k_1 = k_2$ | $[1, 2]$ |
| $m_1 = m_2$ | $[0.1, 0.3]$ |
| $S_1^I, S_2^I$<br>$S_1^F, S_2^F$ | $[0.2, 2]$ |

Similarly we computed the frequency dependence of the community (positive, negative or no frequency dependence) and the superior competitor (species 1, species 2 or equal fitness) for the initial resource supply rates. No-frequency dependence was only ever observed with the species-specific method, which happens for very unbalances resource supply rates, as both species drive one resource to extinction in mono-culture, effectively competing only for one resource. The other methods can lead to no change in niche differences, but the probability for no niche difference is zero. Similarly, we never observed equal fitness of both competitors, which is theoretically possible, but has zero probability of occurring. We again compared how often the different methods agreed and how often we would expect them to agree based on chance.

**Figure S1:**
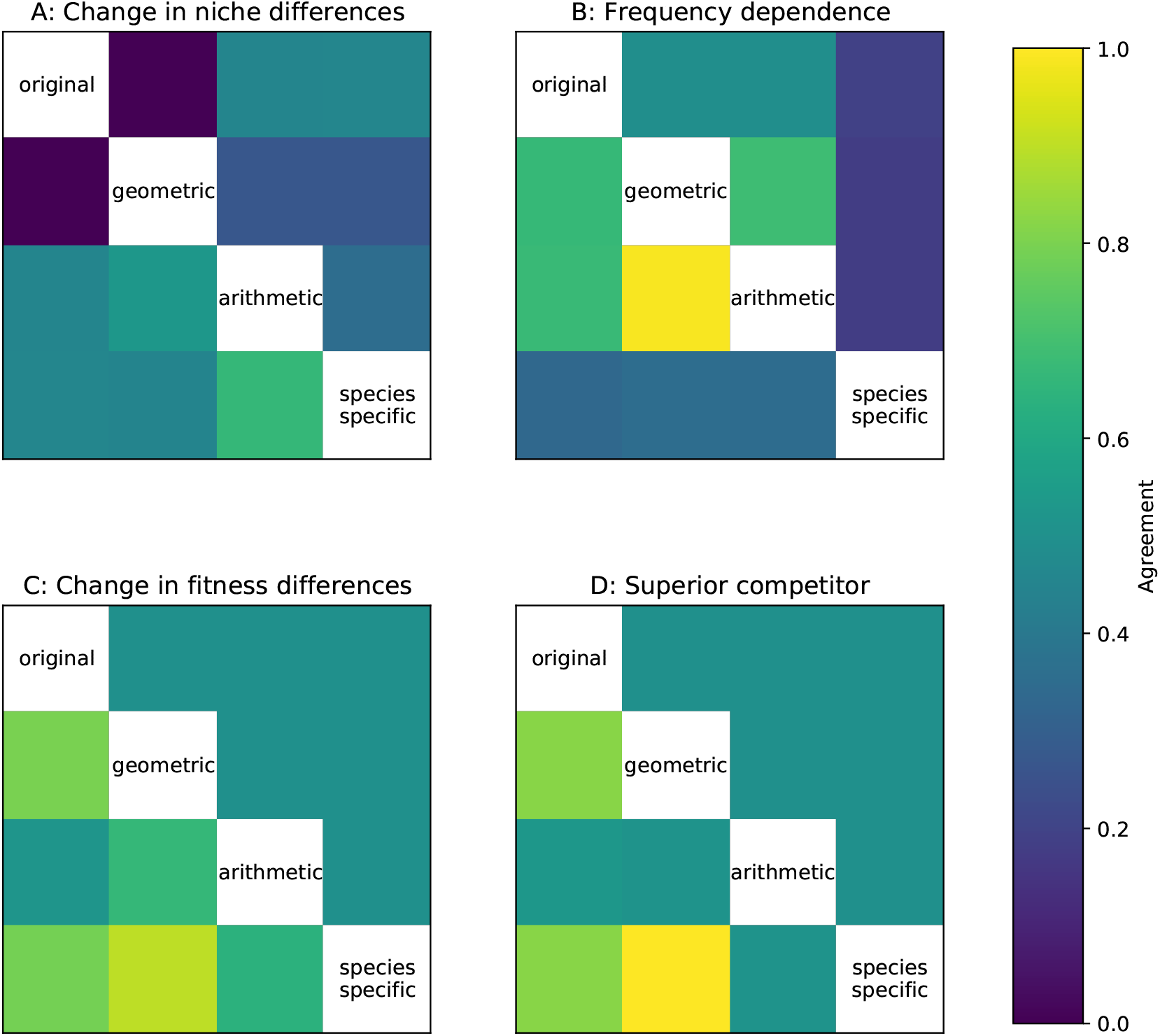
Similar to figure 5. A, C: How often two methods agree on how changes in resource supply rates affect niche differences (A) and fitness differences (C). Entries below diagonal show actual agreement, entries above diagonal show how much we would expect them to agree based on chance. B,D: How often two methods agree on the frequency dependence (B) and the superior competitor (D) for the initial resource supply point. Entries below diagonal show actual agreement, entries above diagonal show how much we would expect them to agree based on chance.

**Figure S2:**
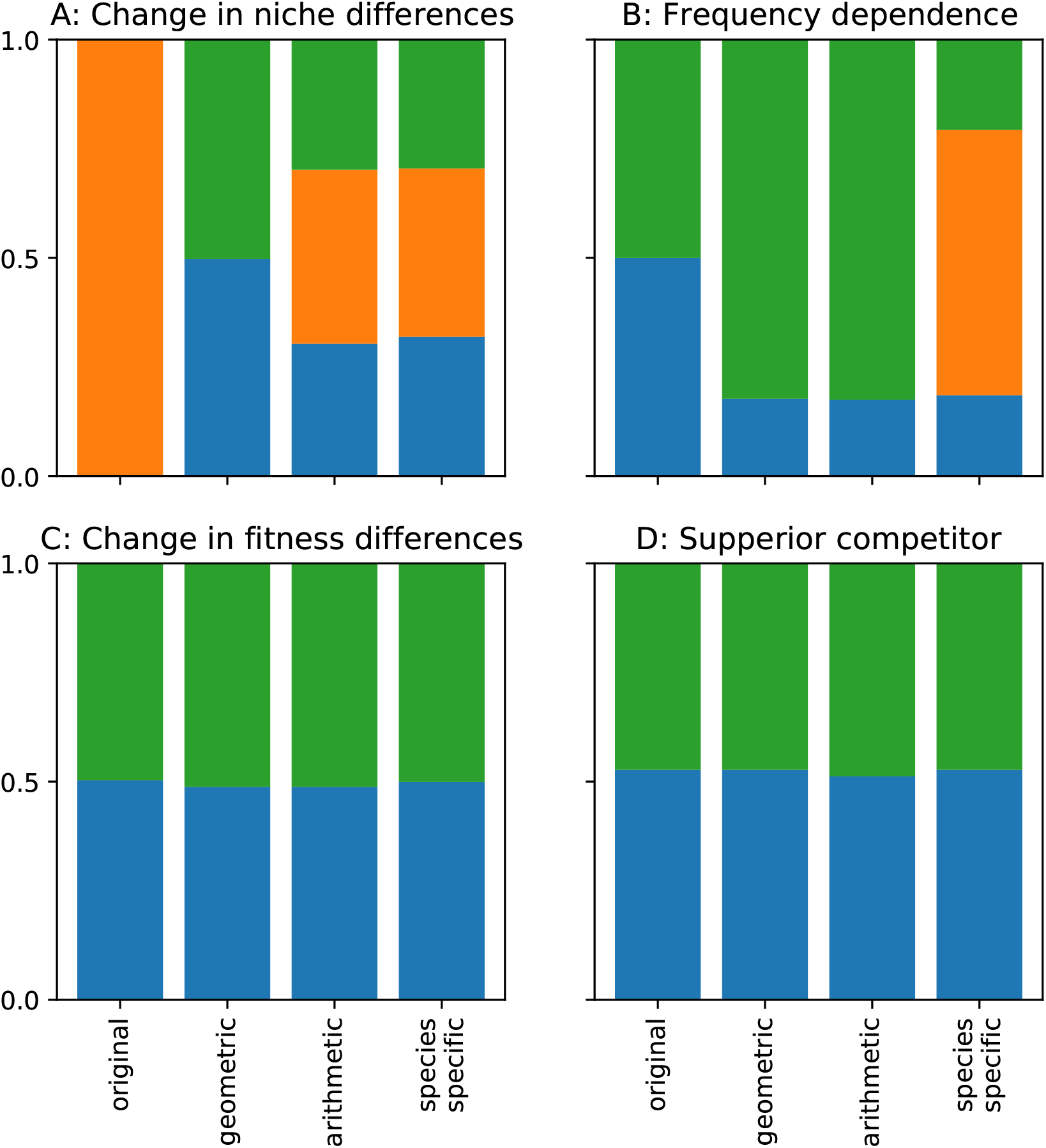
Base probabilities *P*(*x* |*method*) for the different methods for niche and fitness differences investigated for change in niche differences (A), frequency dependence (B), change in fitness differences (C) and superior competitor (D). Colors indicate the three different possible outcomes (blue, orange and green correspond to: A and C: decrease, no change, increase, B: positive,zero and negative, D: species 1, similar fitness and species 2). Panel A corresponds to above diagonal of figure 2B, Panel B to above diagonal 2A, Panel C to below diagonal of Figure 1B, and Panel D to below diagonal of Figure 2A. A: For the original method no changes in resource supply rates affected niche differences, as proven by (Letten *et al*., 2017). For the geometrical method we never observed no-change in niche differences. B: Only the species-specific predicts that some communities have no niche differentiation, this is the case when both species drive one resource to extinction in monoculture and are effectively limited by only on resource. C and D: We never observed that changes in resource supply rates have no effect on fitness differences or that both species have equal fitness. The differences from 0.5 probabilities stem from the randomness of the simulations.

## S2 Growth rates and niche and fitness differences

**Figure S3:**
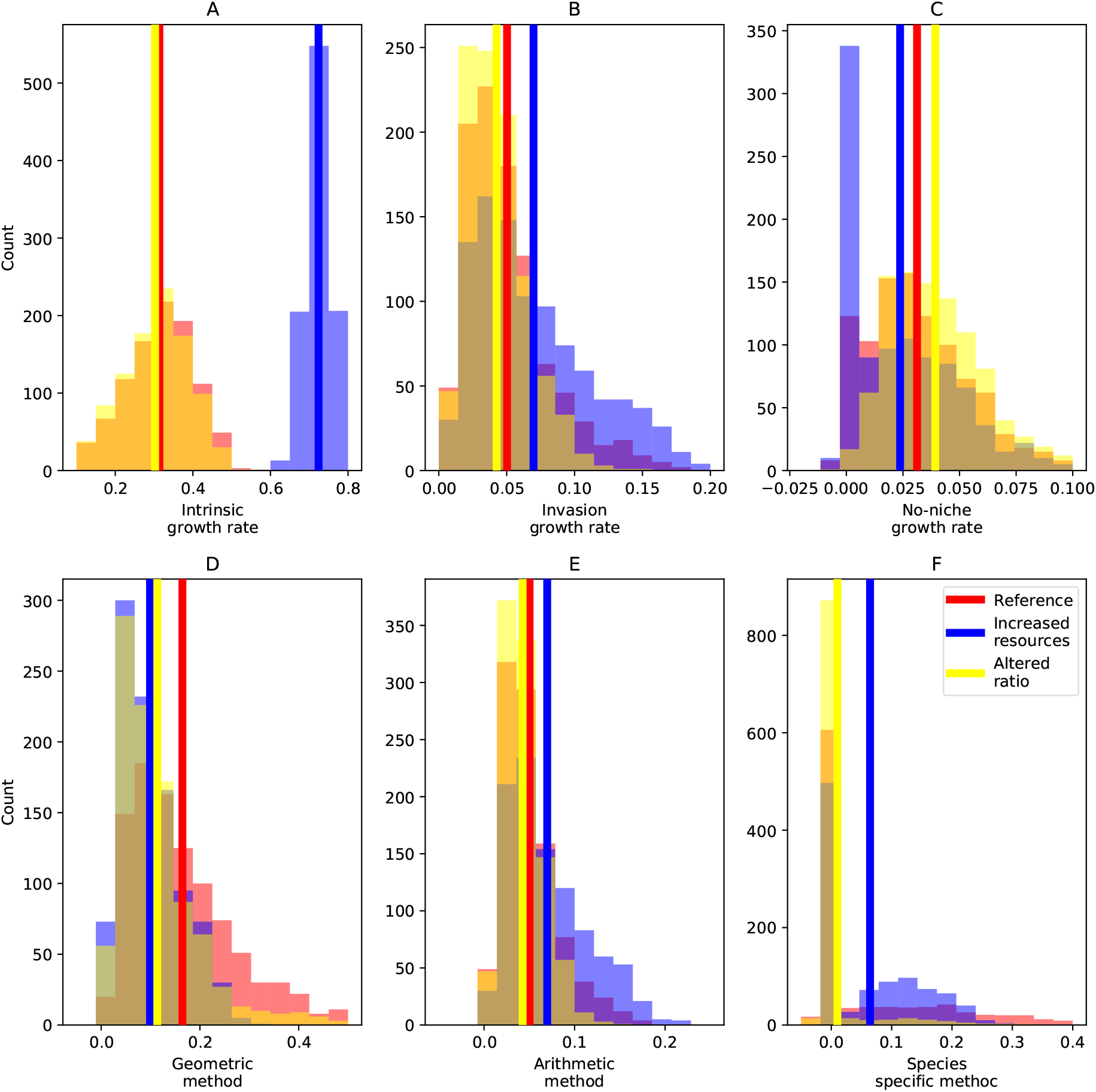
A-C: Histograms of the mean intrinsic (A), invasion (B) and no-niche (C) growth rates for three different resource supply rate scenarios, corresponding to the reference case (red), increased resource supply at fixed ratios (blue) and changed resource supply rates but same total supply rates (yellow). Increasing resource supply rates has a strong positive effect on intrinsic growth rate, a smaller positive effect on invasion growth rates and a negative effect on the no-niche growth rate. Changing resource ratios has no effect on the intrinsic growth rates, but small effects invasion and no-niche growth rates. D-F: Histograms of the niche differences according to the geometric (D), arithmetic (E) and species-specific method (F). Different resource scenarios have different effects on the growth rates and therefore different effects on the niche differences according to the different methods. D: Increasing resource supply rates has a positive effect on both, intrinsic and invasion growth rates. However, the effect on the intrinsic growth rate is stronger, such that niche differences according to the geometric method are decrease with increasing resource supply rates. E: For the arithmetic method, niche differences depend only on the invasion growth rates, such that increasing resources increases niche differences. F: For the species-specific method, niche differences are lowest for the altered resource scenario, as this increases the no-niche growth rates and (slightly) decreases invasion growth rates. A-F: Vertical bars indicate the mean of the corresponding distribution.

**Figure S4:**
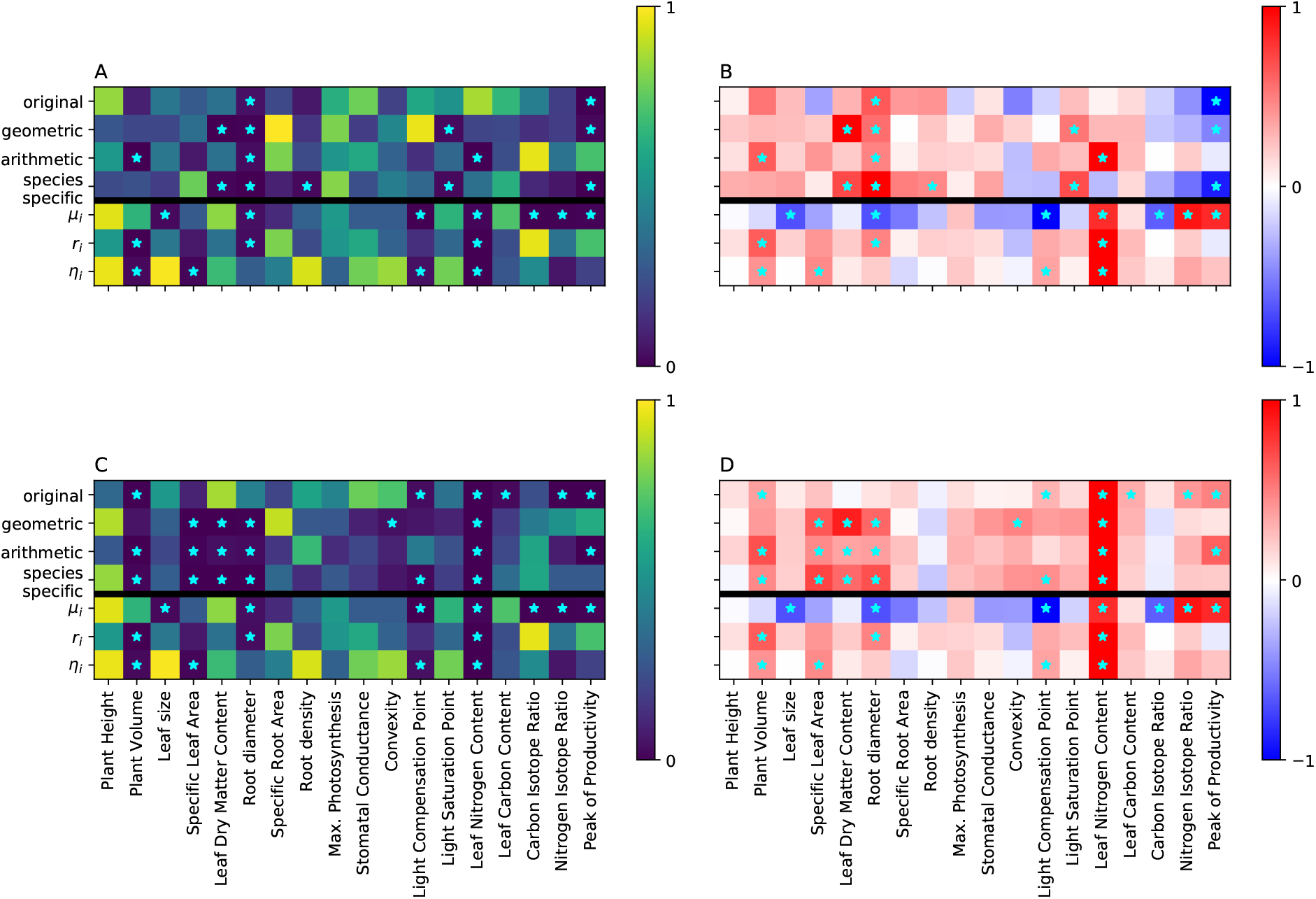
Equivalent to figure 3, the p-value (A,C) and the relative slope (B,D) of the linear regression between the pairwise trait differences (x-axis) and the niche (A,B) and fitness differences (C,D) for each of the four investigated methods (y-axis) for the Pérez-Ramos *et al*. (2019) data set (above the black horizontal line). Below the black horizontal line we show the linear regressions for the mean of the intrinsic growth rate *µ*_*i*_, the invasion growth rate *r*_*i*_ and the no-niche growth rate *η*_*i*_. Intuitively, if a trait correlates with a growth rate, then we would expect that this trait also correlates with niche differences that include this growth rate. For example, the arithmetic method depends only on the invasion growth rate, consequentially, every trait that correlates with the invasion growth rates also correlates with the niche differences as computed by the arithmetic method. However, the different growth rates are also strongly correlated, which complicates the issue. For example, root density does not correlate with any growth rate, yet it does correlate with niche differences according to the species-specific method. Similarly, leaf nitrogen content correlates with all three growth rates, yet it does not correlate with the original, geometric or species-specific method.

## S3 Theoretical comparison of the different methods

In the main text we reported that the different methods lead to different interpretations of niche and fitness differences and coexistence in general. We here give a short insight into where these differences stem from. We first analyse two simple and well known community models (Lotka-Volterra model and Beverteon-Holt annual plant model) and then give some limited insight into the general case without a specific community model.

### S3.1 Lotka-Volterra model

As in the main text, we assume a Lotka-Volterra community model of the form

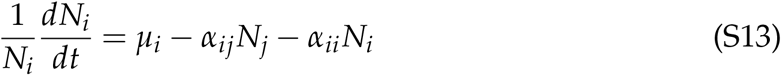

Niche and fitness differences according to the original method are given in equation 1 and 2. Carroll *et al*. (2011) have shown that the geometrical method gives identical values for niche and fitness differences as the original method for the Lotka-Volterra model Similarly, Spaak *et al*. (2020) have shown that the species-specific method gives identical values for niche differences and fitness differences.

The arithmetic method differs from all these methods. To compute niche and fitness differences according to the arithmetic method we have to compute the mono-culture equilibrium densities 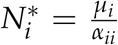 and the invasion growth rates 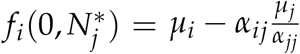 With this we can compute

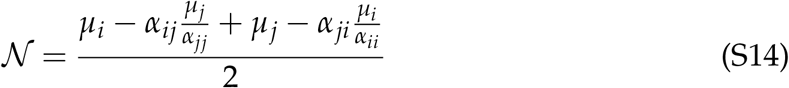

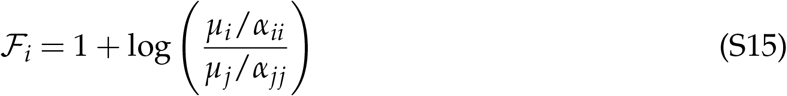

From this we see that the niche by the arithmetic method depend on the intrinsic growth rates *µ*_*i*_ and *µ*_*j*_, while the niche differences as defined by the original method does not, they are therefore qualitatively different. Similarly, the fitness differences by the arithmetic method do not depend on interspecific interaction coefficients *α*_*ij*_ and *α*_*ji*_.

### S3.2 Annual plant model

We here investigate how the different methods compare for the annual plant model, another simple but widely applied community model. For simplicity we assume that the annual plants do not have a seedbank, i.e. we use the following community model:

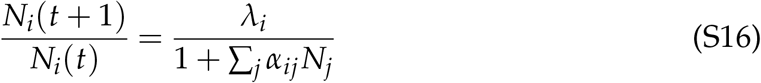

where *N*_*i*_(*t*) is the density of species *i* in year *t, λ*_*i*_ is the basic reproduction rate and *α*_*ij*_ is the per-capita effect of species *j* on species *i*. The annual plant model is time discrete, to apply the niche and fitness differences metrics we first have to convert the growth rates into continuous form by computing the natural logarithm of the growth rates.

The intrinsic growth rates, monoculture growth rates and the invasion growth rates are given by:

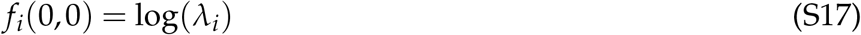

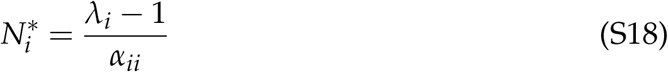

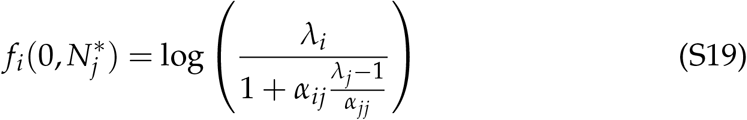

With this we can compute the different methods

For the original method we have

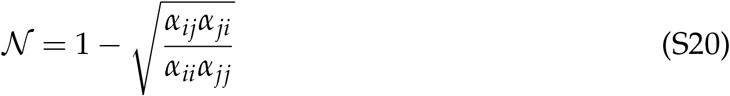

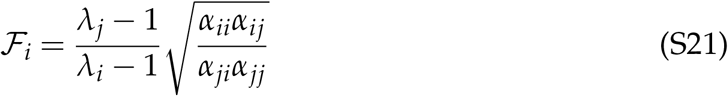

For the geometrical method we have

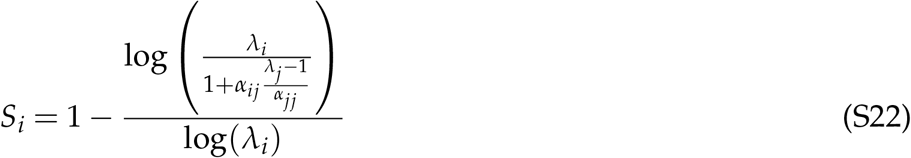

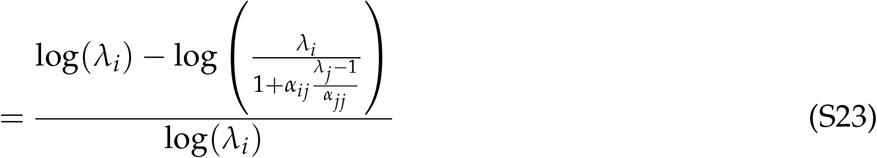

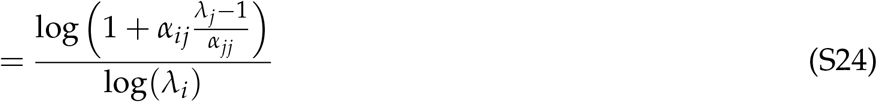

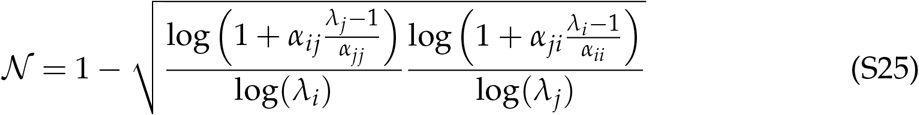

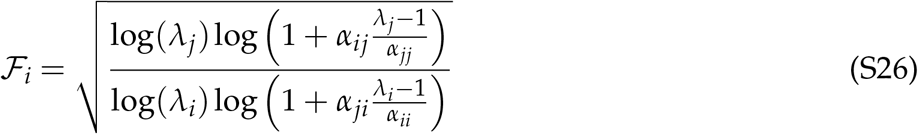

For the arithmetic method we have

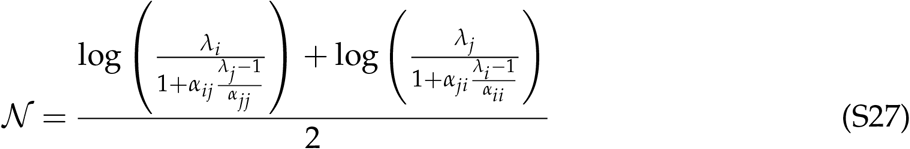

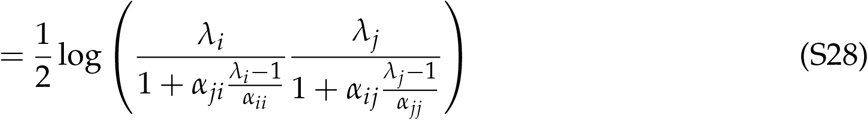

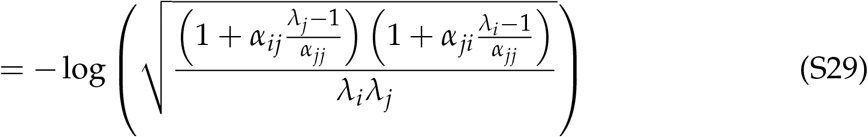

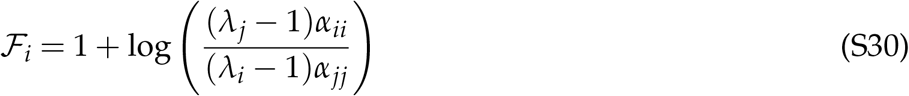

For the species-specific method we have

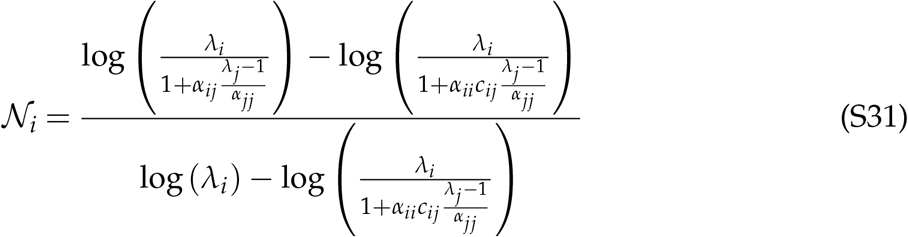

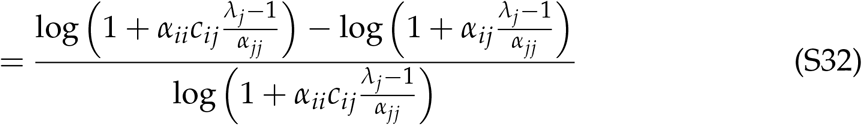

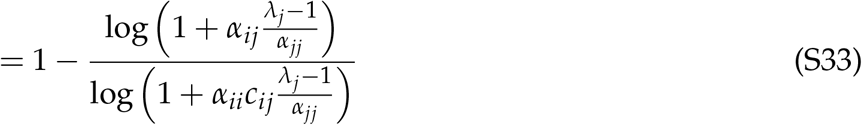

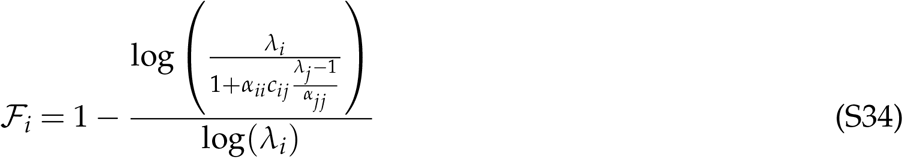

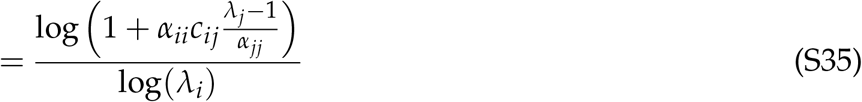

We use 𝒩_*i*_ with subscript *i* in the appendix, as this is needed for clarity and to solve the necessary equations. Unfortunately, we were not able to obtain closed form expressions for the quantities *c*_*ij*_.

These equations show that the mathematical expressions for niche and fitness differences differ for the various methods. However, they give us little intuitive understanding as the equations are quite complex.

Qualitatively, all niche differences depend positively on *α*_*ii*_ and *α*_*jj*_ and negatively on *α*_*ji*_ and *α*_*ij*_. However, their functional responses differ. Conversely, the different methods have a different qualitative dependence on the parameters *λ*_*i*_ and *λ*_*j*_ (Fig. S5). We investigate here the extreme cases *λ*_*i*_, *λ*_*j*_ → 1 and *λ*_*i*_ → ∞. The original method is not affected by change in intrinsic growth rate *λ*_*i*_.

**Figure S5:**
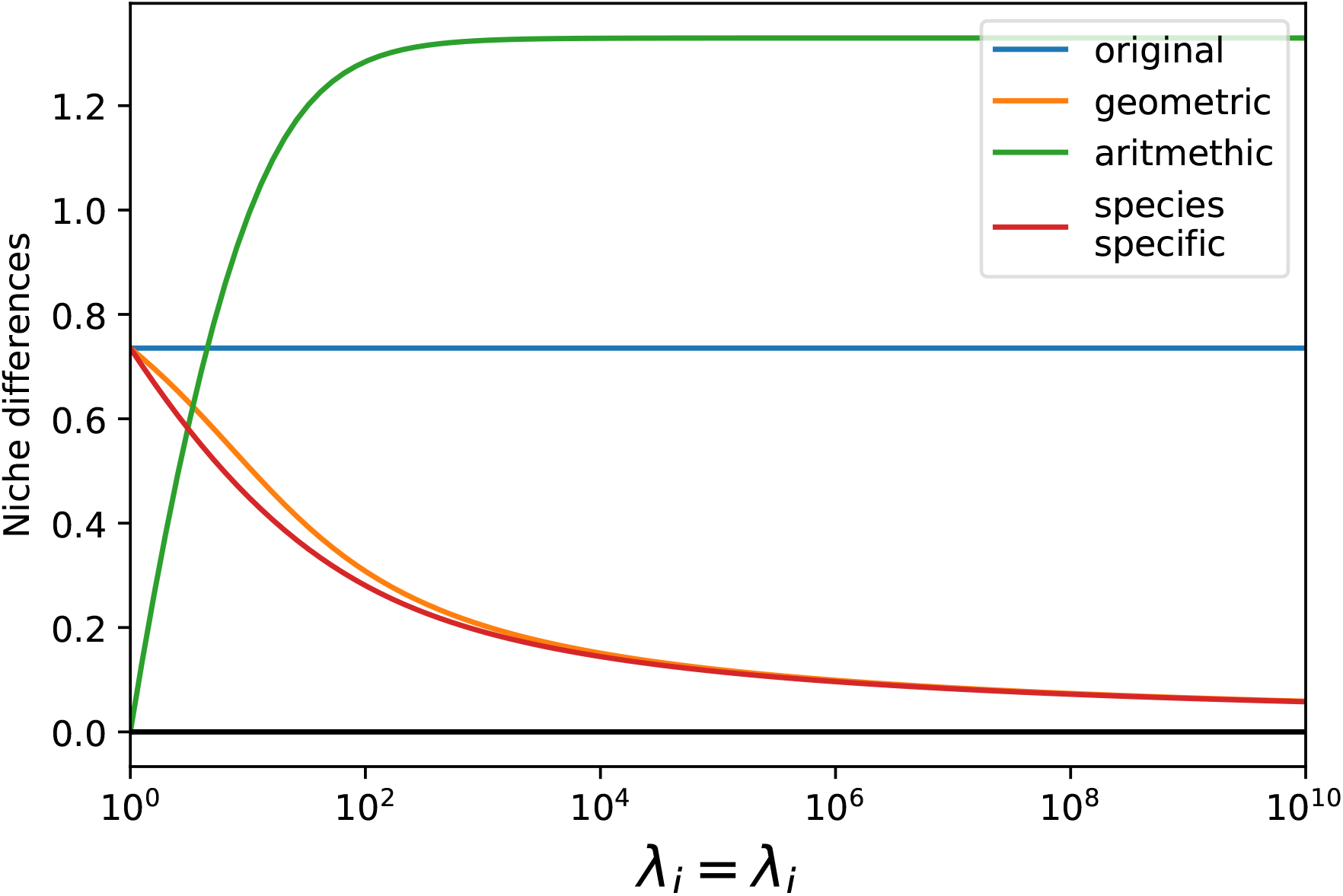
Dependence of niche differences on *λ*_*i*_ for the different methods. Niche differences by the original method are independent of *λ*_*i*_ and *λ*_*j*_. Niche differences by the geometric and species-specific method decrease with increasing *λ*_*i*_ and *λ*_*j*_. In the limit case of *λ*_*i*_ = *λ*_*j*_ = 1 they agree with the original method. In the other limit case of *λ*_*i*_ = *λ*_*j*_ → ∞ they tend to zero. Finally, the arithmetic method increases with increasing *λ*_*i*_.

For the geometric method:

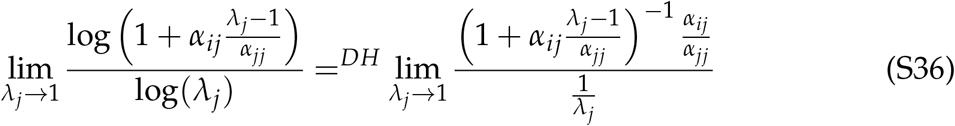

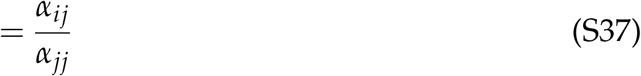

Where =^*DH*^ indicates the de L’Hopital rule. This proves that 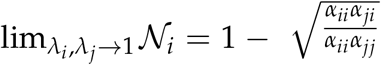*0020*for the geometric method, i.e. in the limit case the geometric and the original method are identical. For the other extreme case:

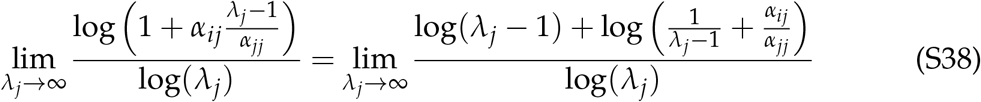

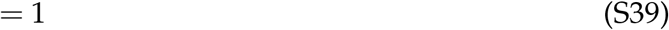

**Table S2:**
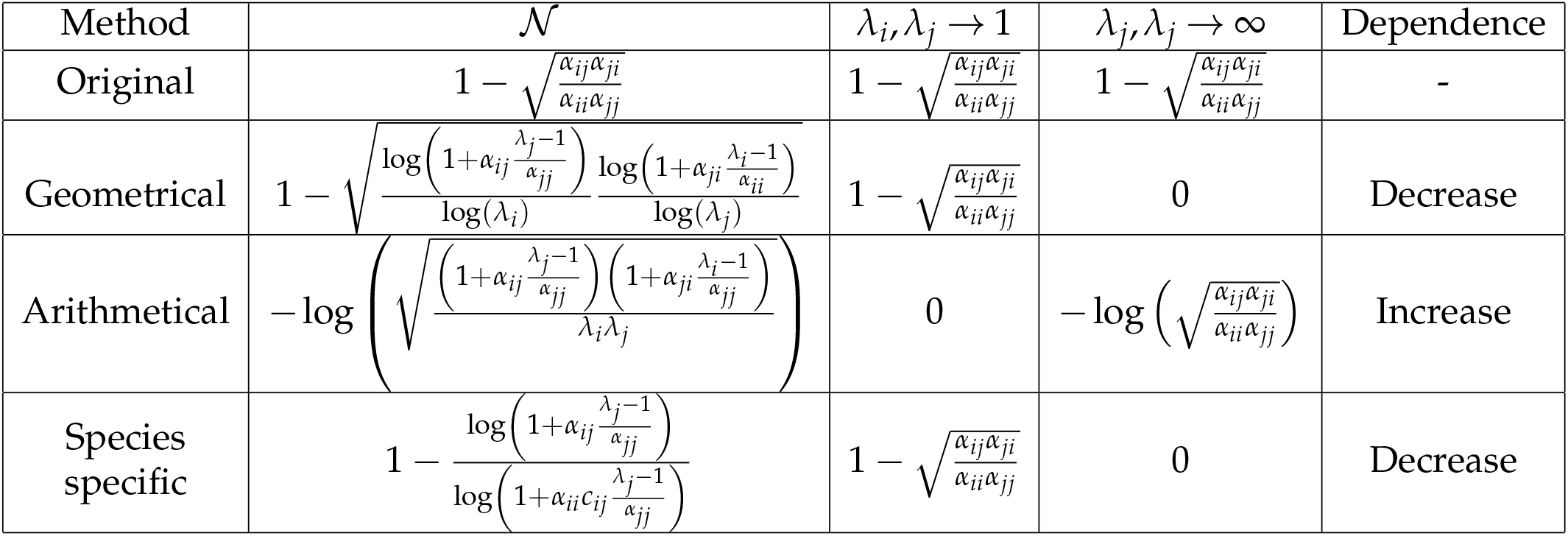
Niche and fitness differences according to the different methods for the annual plant method. gives the mathematical expression for the niche differences. *λ*_*i*_, *λ*_*j*_ → 1 gives the limit case for when the intrinsic growth rates approach 1. *λ*_*i*_, *λ*_*j*_→∞ gives the limit case for when the intrinsic growth rates approach ∞. Dependence:describes how 𝒩_*i*_ depends on the intrinsic growth rates for the case of 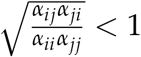. However, this dependence might not be monotone.

Which proves that 𝒩 → 0 for increasing *λ*_*i*_, *λ*_*j*_ → ∞. In general, increasing intrinsic growth *λ*_*i*_, *λ*_*j*_ therefore bring niche differences closer to 0 in the geometric method.

For the arithmetic method:

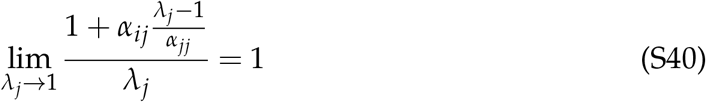

which gives 𝒩 → 0 for the arithmetic method. Conversely,

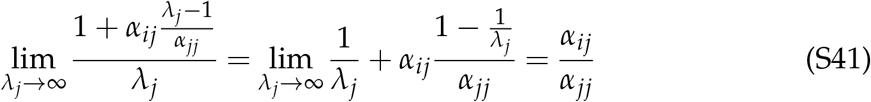

Which gives 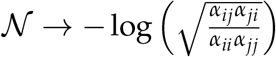 for increasing *λ*_*i*_, *λ*_*j*_ → ∞. In general, decreasing intrinsic growth *λ*_*i*_, *λ*_*j*_ therefore bring niche differences closer to 0 in the arithmetic method, unlike in the geometric method.

For the species-specific method

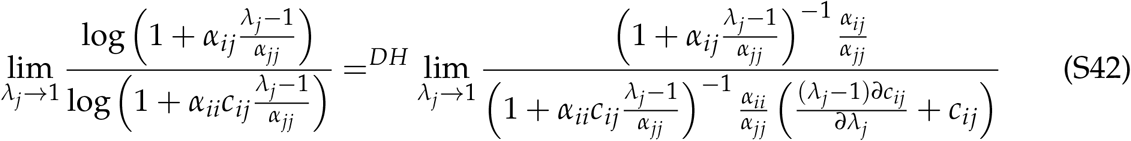

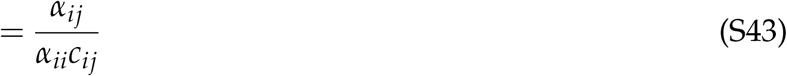

Similarly, we get 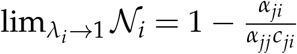. By definition we have 𝒩_*i*_ = 𝒩_*j*_ which yields 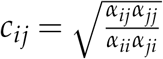 and 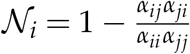.

For increasing *λ*_*j*_ we have:

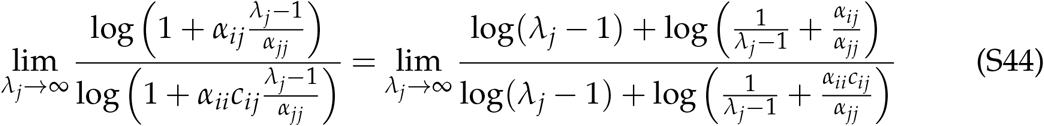

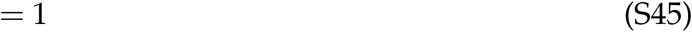

Which yields 𝒩_*i*_ → 0 for *λ*_*j*_ → ∞. Consequently, similar to the geometric framework the species-specific framework approaches the original method for decreasing *λ*_*i*_, *λ*_*j*_. For increasing *λ*_*j*_, 𝒩_*i*_ approaches 0. However, for the geometric method both *λ*_*i*_ and *λ*_*j*_ must approach ∞ for zero niche differences, while for the species-specific method only one of the two has to approach ∞.

### S3.3 Model independent

Unfortunately, little can be said for a general community model. We focus here on the arithmetic, geometric and species-specific method, as these can be applied to any community model.

Qualitatively the methods agree on the sign of niche differences for a community driven by coexistence or priority effects. For a community driven by coexistence we have 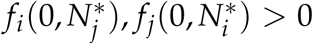, which leads to *S*_*i*_, *S*_*j*_ < 1. Consequently, we have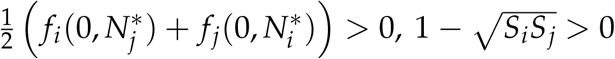. Finally, Spaak *et al*. (2020) have shown coexisting species are driven by negative frequency dependence and therefore have positive niche differences. The case of priority effects is equivalent with all ineqaulity signs exchanged. These three methods therefore qualitatively disagree only for the case of competitive exclusion.

Next, we show that the species-specific and geometric method always agree on the competitive superior species. We assume without loss of generality that species *i* is the superior competitor by the species-specific method, we therefore have

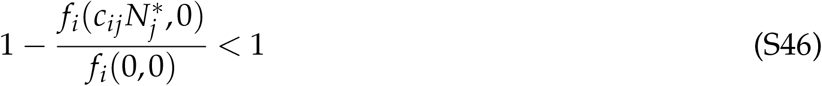

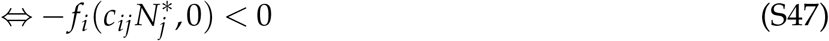

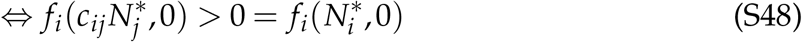

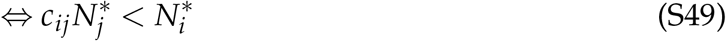

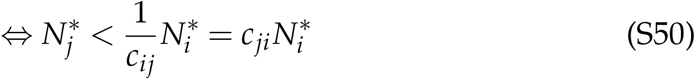

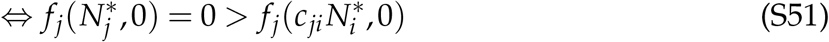

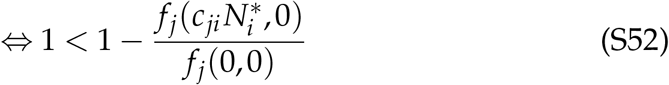

Where we used the monotony of the per-capita growth rates from S48 to S49 and from S50 to S51, and we used the definition of 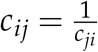 from S49 to S50. We therefore indeed have only one species with a fitness difference exceeding 1. And only for this species (by assumption species *i*) the no-niche growth rate 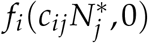 is positive.

Rearranging this we get

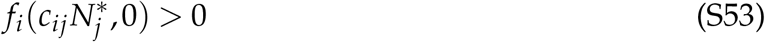

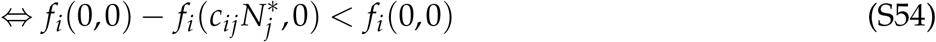

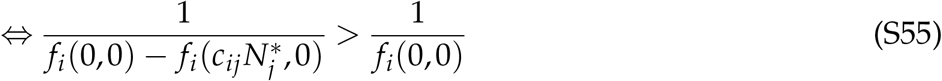

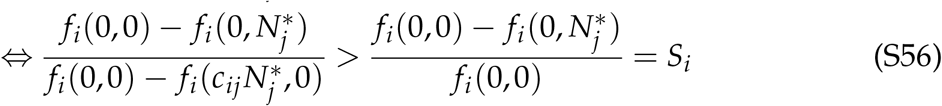

where the left hand side of S56 is the niche overlap as defined by the species-specific method and the right hand side of S56 is the sensitivity of species *i* as defined by the method of Carroll. As species *i* was the competitive dominant by the species-specific method we therefore must have

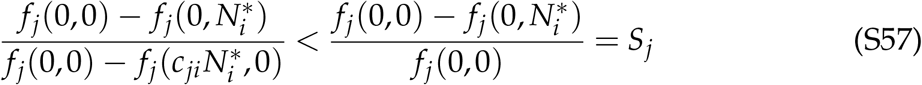

Additionally, we have by the definition of the species-specific method

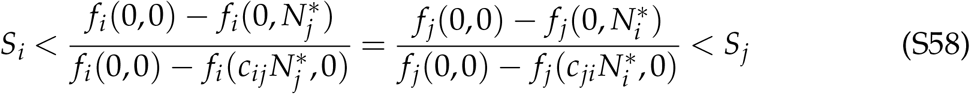

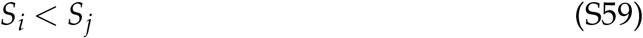

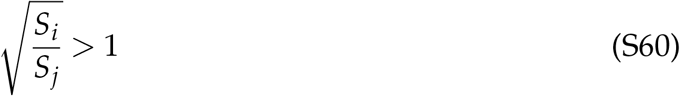

Therefore the species-specific and the geometric method always agree on the competitive dominant species.

## S4 Quantitative comparison for the annual plant data

**Figure S6:**
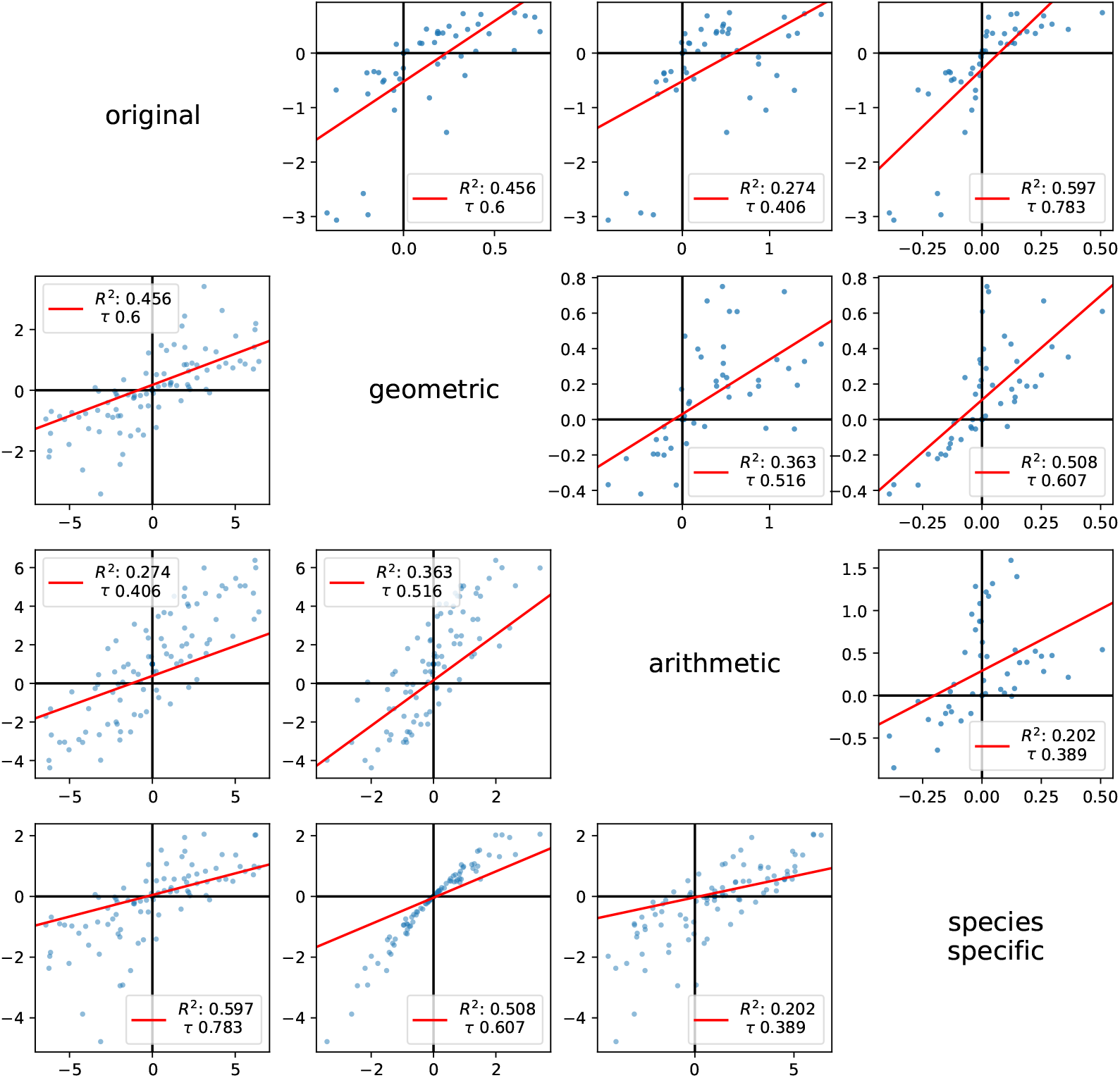
In the main text we focused on the qualitative differences between the methods. Here we show a more quantitative comparison for the data from Pérez-Ramos *et al*. (2019). We plot the niche (panels above diagonal) and fitness differences (panels below diagonal) of each species by each method and compare them. The fitness differences have been log-transformed. For example, the top-right plot shows the niche differences as computed by the species-specific method on y-axis and the niche differences as computed by the arithmetic method on the x-axis. Similarly, for the other plots. We performed a linear regression (red line) and report it’s *R*2 value. Some relationships were not linear, so we also report the Kendal-Tau correlation for non-parametric regression.

**Figure S7:**
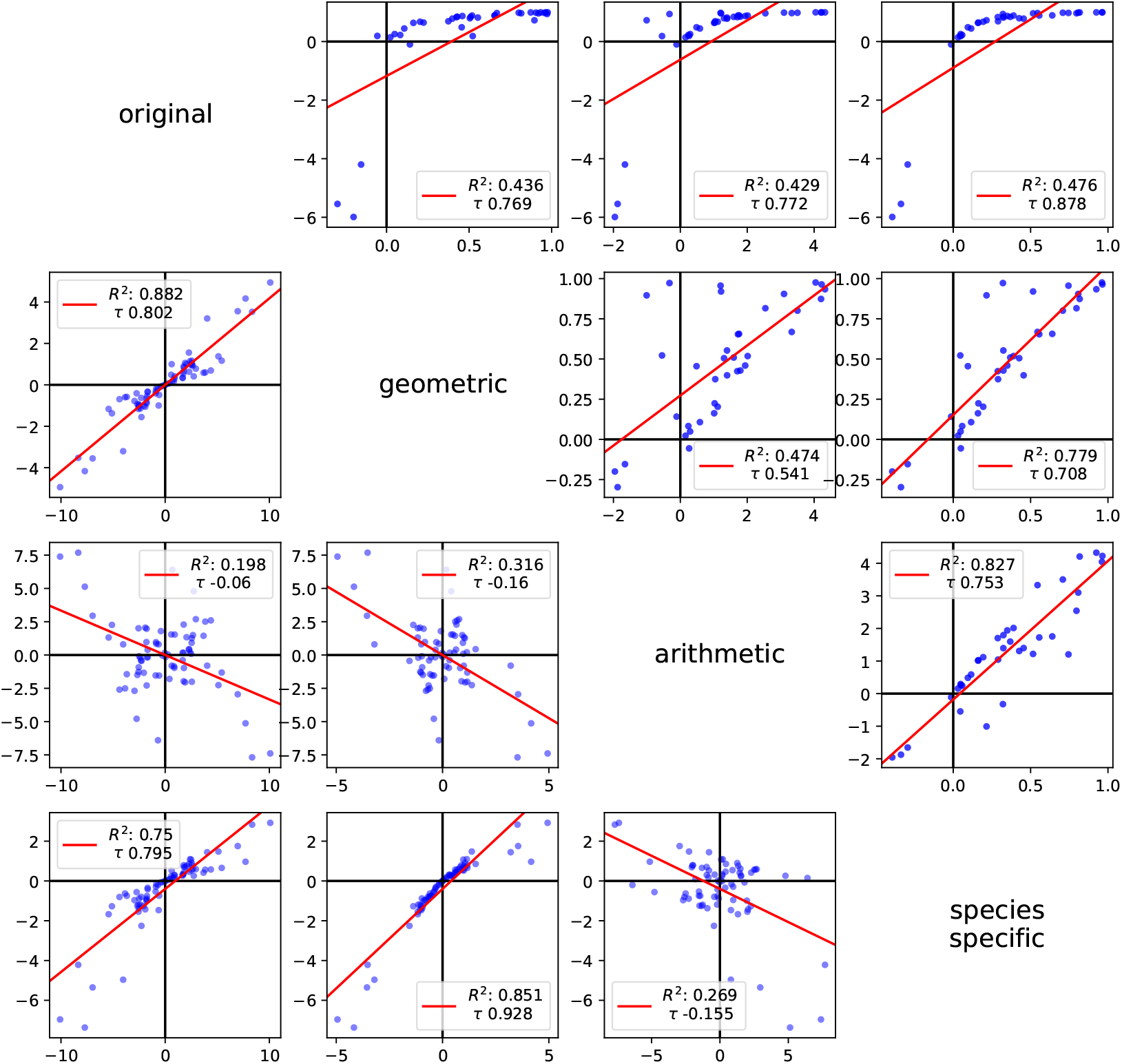
Similar to figure S6 but for the data from Germain *et al*. (2016).

## Notes

### Competing Interest Statement

The authors have declared no competing interest.

